# CD55 regulates self-renewal and cisplatin resistance in endometrioid tumors

**DOI:** 10.1101/155887

**Authors:** Caner Saygin, Andrew Wiechert, Vinay S. Rao, Ravi Alluri, Elizabeth Connor, Praveena S. Thiagarajan, James S. Hale, Yan Li, Anastasia Chumakova, Awad Jarrar, Yvonne Parker, Daniel J Lindner, Anil Belur Nagaraj, J Julie Kim, Analisa DiFeo, Fadi W. Abdul-Karim, Chad Michener, Peter G. Rose, Robert DeBernardo, Haider Mahdi, Keith R. McCrae, Feng Lin, Justin D. Lathia, Ofer Reizes

**Author notes:** These senior authors contributed equally (Justin D. Lathia, Ofer Reizes). **Contact:** Dr. Justin D. Lathia, Department of Cellular and Molecular Medicine, Lerner Research Institute, 9500 Euclid Ave., NC10, Cleveland, OH 44195, USA; Dr. Ofer Reizes, Department of Cellular and Molecular Medicine, Lerner Research Institute, 9500 Euclid Ave., NC10, Cleveland, OH 44195, USA.

## Abstract

Effective targeting of cancer stem cells (CSCs) requires neutralization of self-renewal and chemoresistance, however these phenotypes are often regulated by distinct molecular mechanisms. Here we report the ability to target both of these phenotypes via CD55, an intrinsic cell surface complement inhibitor, which was identified in a comparative analysis between CSCs and non-CSCs in endometrioid cancer models. In this context, CD55 functions in a complement-independent manner and required lipid raft localization for CSC maintenance and cisplatin resistance. CD55 regulated self-renewal and core pluripotency genes via ROR2/JNK signaling and in parallel cisplatin resistance via LCK signaling, which induced DNA repair genes. Targeting LCK signaling via saracatinib, an inhibitor currently undergoing clinical evaluation, sensitized chemoresistant cells to cisplatin. Collectively, our findings identify CD55 as a unique signaling node that drives self-renewal and therapeutic resistance via a bifurcating signaling axis and provide an opportunity to target both signaling pathways in endometrioid tumors.

**SUMMARY:** CD55 is a membrane complement regulatory protein that attenuates complement-mediated cytotoxicity. Saygin et al. elucidate a new role for CD55 as a signaling hub for cancer stem cell self-renewal and cisplatin resistance pathways in endometrioid tumors and open a new line of research into chemotherapeutic-refractory cancers.

**Abbreviations:**

GPI: glycophosphatidylinositol
LIME: LCK interacting transmembrane adaptor
PAG: protein associated with glycosphingolipid-enriched microdomains
CSC: cancer stem cell
DAF: decay accelerating protein
LCK: Endometrioid Tumor, ET
CSC: cancer stem cell
mCRP: membrane-bound complement regulatory protein
ROR2: receptor tyrosine kinase like orphan receptor 2
LCK: lymphocyte-specific protein tyrosine kinase
JNK: c-Jun N-terminal kinase

## INTRODUCTION

Uterine and ovarian cancers are the most common gynecological cancers in the US (Baldwin LA HB, 2012; Siegel RL, 2016). These tumors are characterized by four main histological subtypes: endometrioid, serous, mucinous, and clear cell carcinoma (Karst AM, 2010; Kurman RJ, 2016). Endometrioid carcinomas make up over 80% of uterine cancers and contribute to 15% of epithelial ovarian cancers (DiSaia PJ, 2012). Endometrioid uterine and ovarian cancers are thought to arise from similar cells of origin (Catasus L, 2009; Cuellar-Partida G, 2016). In advanced stage disease, both uterine and ovarian cancers are treated with cytoreductive surgery and platinum-based cytotoxic chemotherapy (Armstrong DK BB, 2006). While many patients achieve clinical remission with this standard approach, advanced stage uterine and ovarian cancers are prone to recurrence (Hanker LC, 2012). Chemoresistance is generally defined as progression of disease within 6 months of therapy. Patients with relapsed disease are considered incurable in most cases and management is intended to prolong life with symptomatic relief (Hanker LC, 2012). Several genomic studies have demonstrated that endometrioid tumors are genetically heterogeneous with diverse molecular subtypes, and an actionable driver gene mutation has not been identified (Cancer Genome Atlas Research Network., 2013; CGAR, 2011; Tan TZ, 2013). Therefore, there is an increasing need to identify pathways driving cisplatin resistance that can be targeted to overcome resistance, which otherwise presents as incurable disease.

Both uterine and ovarian endometrioid tumors are heterogeneous and shown to contain a self-renewing cancer stem cell (CSC) population. CSCs are implicated in tumor recurrence and treatment resistance (Kyo S, 2011; Nagaraj AB, 2015; Wiechert A, 2016). Endometrioid CSCs can be isolated by well-established surface markers, including CD133, CD44, CD49f, ALDH activity, and stem cell reporter systems (Kyo S, 2011; Wiechert A, 2016). Utilizing multiple CSC enrichment methods, we identified that decay accelerating factor (CD55) was highly expressed in endometrioid CSCs and cisplatin-resistant cells. CD55 is a glycophosphatidylinositol (GPI)-anchored membrane complement regulatory protein (mCRP) that protects cells from complement-mediated lysis (Lukacik P, 2004). It is shown to be expressed in ovarian and uterine cancers, and the levels are higher in malignant vs benign endometrial tissue (Kapka-Skrzypczak L, 2015; Murray KP, 2000). CD55 expression was also shown to have a prognostic significance in patients with breast cancer (Ikeda J, 2008). In addition to the canonical effects including the modulation of the efficiency of anti-tumor monoclonal antibodies, CD55 has been shown to signal intracellularly and activate receptor tyrosine kinases at lipid rafts (Shenoy-Scaria AM, 1992). The role of non-canonical CD55 signaling in T cell receptor activation has been well characterized, however there are limited studies on the intracellular actions of CD55 in cancer (Ventimiglia LN, 2013). Based on our initial findings in complement-free conditions, we hypothesized that CD55 may regulate self-renewal and cisplatin resistance in endometrioid tumors via a complement-independent mechanism.

## RESULTS

### CD55 is highly expressed in CSCs and cisplatin resistant cells

We have recently validated the NANOG promoter-driven green fluorescence protein (GFP) reporter system in isolation of endometrioid CSCs (Wiechert A, 2016). We used NANOG-GFP reporter-transduced cisplatin-naive (A2780) and-resistant (CP70) ovarian endometrioid tumor cell lines to perform a high throughput flow cytometry screen **(Fig. 1A)**. Out of 242 cell surface markers included in the screening panel, CD55 was the most differentially expressed protein in between A2780 CSCs (GFP+) and non-CSCs (GFP-) **(Fig. 1B)**. Both GFP+ and GFP- CP70 cells had high levels of CD55 expression, which might be attributed to the higher self-renewal potential and stem-like properties in cisplatin resistant cells (Wiechert A, 2016). Of the other two mCRPs included in the screen, CD59 was expressed higher in CSCs, while there was no appreciable difference in CD46 expression **(Supplementary Fig. S1A)**. We have further validated these results in several cisplatin-naive endometrioid tumor cell lines (A2780, TOV112D) and a patient-derived xenograft (EEC-4), at the protein and RNA levels **(Fig. 1C-D; Supplementary Fig. S1B-D)**. Moreover, higher CD55 expression was observed in CSCs of two primary uterine endometrioid tumor specimens (UTE-1 and UTE-2) **(Supplementary Fig. S1E)**. In addition, cisplatin resistant (CP70) cells had higher expression of CD55 and CD59 at protein and RNA levels, as compared to their isogenic cisplatin-naive (A2780) counterparts **(Fig. 1E)**. CP70 cells had 186 and 4 fold higher expression of CD55 and CD59 mRNA as compared to A2780 cells, respectively **(Fig. 1E)**. We previously reported that CD49f can enrich a self-renewing population in cisplatin-resistant cells (Wiechert A, 2016). Using this marker, CSCs (CD49f+) isolated from cisplatin resistant ovarian (CP70) and endometrial (HEC1a) cells had higher levels of CD55 as compared to non-CSCs (CD49f-) **(Supplementary Fig. S1F-G)**. To assess CD55 as a marker of CSCs, we performed limiting dilution sphere formation analysis that provides readout for self-renewal, proliferation, and survival. We found that CD55+ cells isolated from cisplatin-naive (A2780, TOV112D, PDX) and-resistant (CP70, HEC1a) endometrioid tumor cells were significantly more self-renewing than their CD55- counterparts (stem cell frequencies for CD55+ vs CD55- were 1 in 2.2 vs 1 in 4.3 in A2780 [p<0.01], 1 in 10.8 vs 1 in 59.2 in TOV112D [p<0.001], 1 in 36 vs 1 in 87.7 in PDX [p<0.05], 1 in 1.4 vs 1 in 5.1 in CP70 [p<0.001], 1 in 59.6 vs 1 in 209.7 in HEC1a [p<0.01]) **(Fig. 1F, Supplementary Fig. 1H)**. We next investigated the utility of CD55 in predicting outcomes of patients with endometrioid ovarian cancer by using K-M plotter biomarker assessment database(Gyorffy B, 2012). Patients with high tumor CD55 expression at diagnosis had significantly worse progression-free survival, as compared to patients with low CD55 levels (hazard ratio= 4.7, confidence interval= 1.5-14.6, p=0.003) **(Fig. 1G)**. These data demonstrate that CD55 is highly expressed in endometrioid CSCs and cisplatin-resistant cells, enriched in self-renewing populations in both cisplatin-naive and -resistant tumors, and predicts survival in patients with endometrioid tumors.

**Figure 1.**
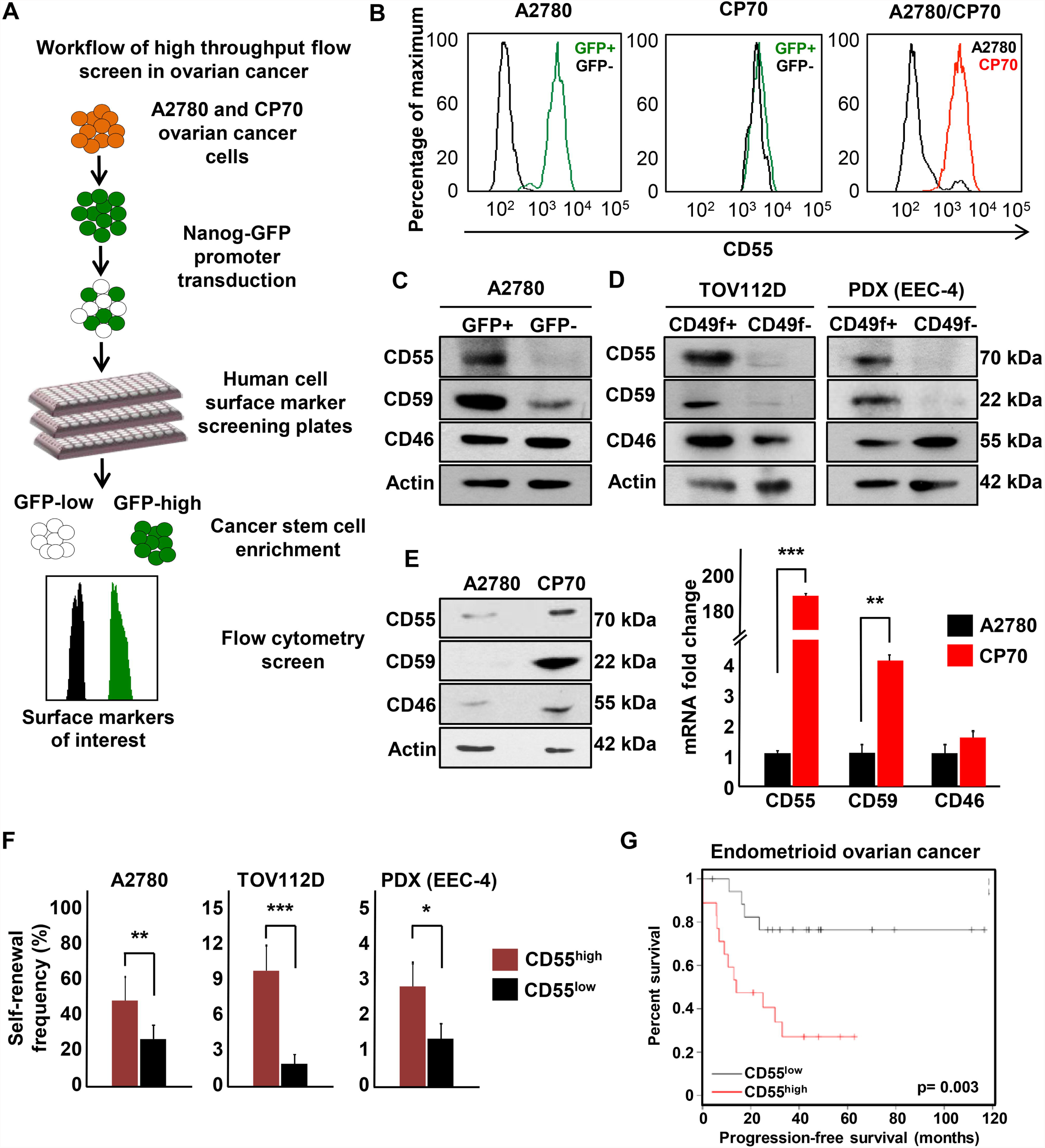
CD55 is highly expressed on endometrioid ovarian and uterine CSCs, and cisplatin-resistant cells. **(A)** A high-throughput flow cytometry screen of 242 different surface CD markers in cisplatin-naïve (A2780) and –resistant (CP70) ovarian cancer cells was performed to investigate the differential expression of these markers between CSCs vs non-CSCs, and cisplatin-naïve vs resistant cells. **(B)** Out of 242 markers, CD55 was the most highly and differentially expressed between cisplatin-naïve CSCs vs non-CSCs, and cisplatin-resistant vs –naïve cells. **(C, D)** Cell lysates from cisplatin-naïve A2780 reporter, TOV112D, and PDX (EEC-4) cells sorted into CSCs and non-CSCs by GFP expression and CD49f expression, respectively, were probed with anti-CD55, CD59, and CD46 antibodies. Actin was used as a loading control. <u>Data are representative of three independent experiments.</u> **(E)** Protein and mRNA expression of CD55, CD59, and CD46 were assessed in lysates from cisplatin-naïve (A2780) and resistant (CP70) cells. Actin was used as a control. <u>Data are representative of two independent experiments.</u> **(F)** Limiting dilution analysis plots of CD55+ compared with CD55- cisplatin-naïve cells. The graph represents the estimates in percentage of self-renewal frequency in sorted populations with the corresponding p-values. <u>Data represent two independent experiments.</u> **(G)** Kaplan-Meier (K-M) progression-free survival curve for endometrioid ovarian cancer patients who had high vs low tumor CD55 expression prior to therapy was obtained from K-M plotter database (<u>http://kmplot.com/analysis/</u>). *p< 0.5, **p< 0.01, ***p< 0.001

### CD55 is necessary for maintenance of stemness and cisplatin resistance

To investigate functional impact of CD55 in CSCs and cisplatin resistant cells, we utilized a genetic approach to inhibit CD55 expression. Using two non-overlapping CD55 shRNA silencing constructs, we inhibited CD55 mRNA and protein expression in both CSCs and cisplatin-resistant cells, but did not impact the expression of CD46 or CD59 **(Fig. 2A, Supplementary Fig. S2A-C)**. Upon CD55 inhibition, core pluripotency transcription factors (NANOG, SOX2, and OCT4) expression was inhibited at the RNA and protein levels **(Fig. 2A, Supplemantary Fig. S2B-C)**. Concomitantly, we observed a decrease in GFP signal intensity in A2780 CSCs, which indicated decreased NANOG promoter activity **(Fig. 2B)**. Limiting dilution tumor sphere formation analysis demonstrated that upon CD55 silencing, cisplatin-naive CSC cultures (A2780, TOV112D, PDX) and cisplatin resistant parental cell cultures (CP70, HEC1a) showed significantly lower self-renewal and stem cell frequencies (reduced from non-targeted control to CD55 knockdown conditions as 1 in 4.8 to 1 in 14.6 and 1 in 10.6 for A2780 CSCs [p<0.001]; 1 in 18.6 to 1 in 41.5 [p<0.01] and 1 in 65 [p<0.001] for TOV112D CSCs; 1 in 21.9 to 1 in 100 and 1 in 207.1 for PDX CSCs [p<0.001]; 1 in 3.3 to 1 in 9.6 [p<0.001] and 1 in 5.9 [p<0.01] for CP70 parental; 1 in 22 to 1 in 50.2 [p<0.01] and 1 in 89.1 [p<0.001] for HEC1a parental) **(Fig. 2C, Supplementary Fig. S2D)**. Since the gold standard functional CSC assay is limiting dilution tumor initiation *in vivo*, we injected CD55-silenced and non-targeted control CSCs into immune-compromised mice at 10^3^, 10^4^, and 10^5^ cells per mouse **(Fig. 2D)**. CD55 silenced cells initiated tumors at a frequency of 1 in 78,398 with the first shRNA construct (p< 0.001), and none of the mice injected with the second construct developed tumors (p< 0.001) compared to a frequency of 1 in 4,522 in non-targeted cells **(Fig. 2D)**. These data provide evidence that CD55 is necessary for CSC maintenance and tumor initiation.

**Figure 2.**
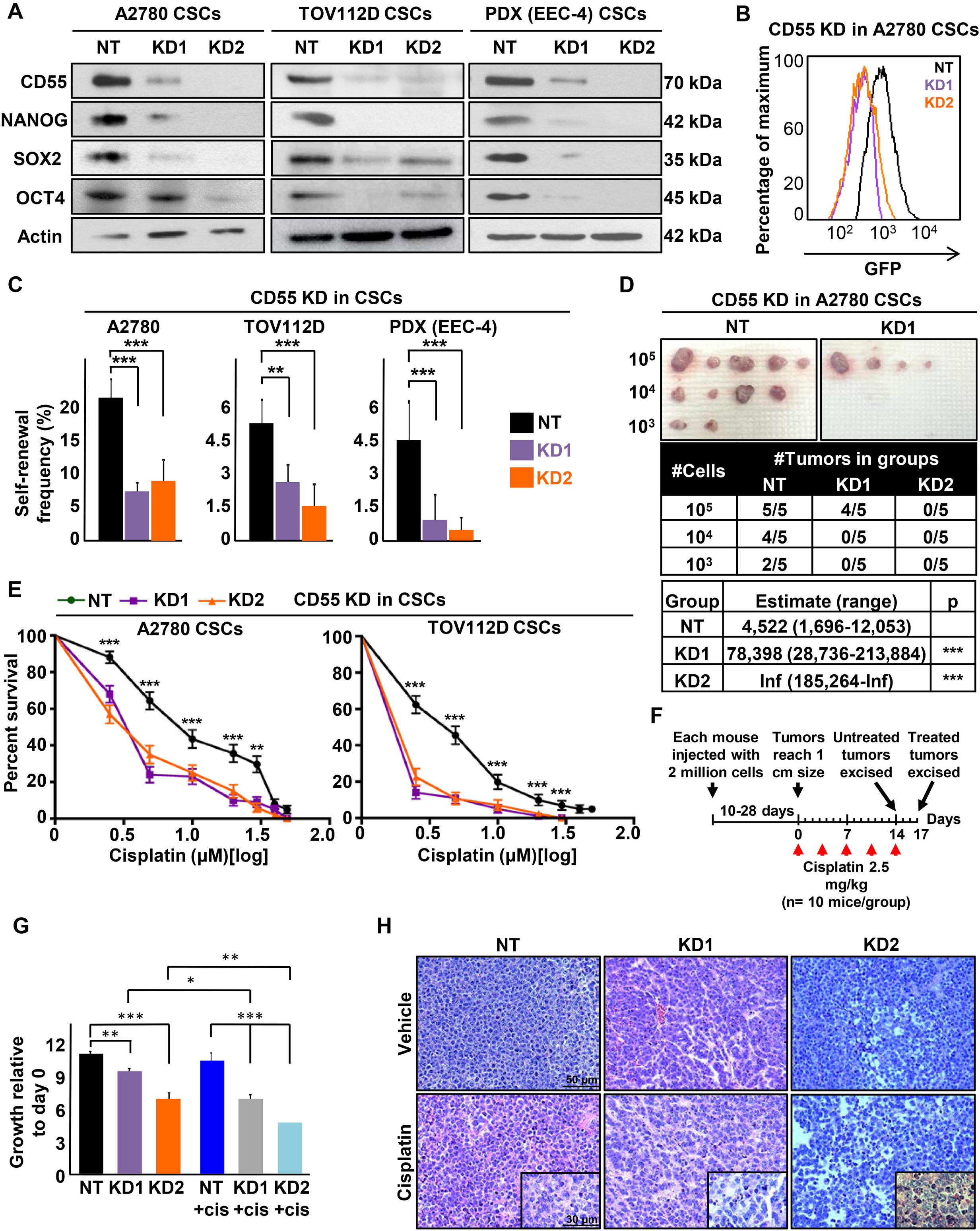
CD55 maintains self-renewal and cisplatin resistance in endometrioid tumors. **(A)** Cell lysates from cisplatin-naïve CSCs silenced for CD55 using two CD55 shRNA constructs (KD1, KD2) and a non-targeting shRNA (NT) control were probed for CD55, NANOG, SOX2, and OCT4 antibodies. Actin was used as a loading control. <u>Data are representative of two to three independent experiments.</u> **(B)** A2780 CSCs silenced for CD55 and NT controls were flowed for GFP signal intensity, which indicates *NANOG* promoter activity. **(C)** Limiting dilution analysis plots of CD55 NT control compared with CD55 KD1 and KD2 silencing constructs in cisplatin-naïve CSCs. **(D)** *In vivo* tumor initiation studies were performed with five mice per group, and the estimates of stem cell frequencies of CD55 NT control compared with the CD55 KD1 and KD2 silencing constructs are shown. **(E)** CD55 silenced cisplatin-naïve CSCs and their NT controls were treated with 0-50 μM cisplatin and percent surviving cells are graphed. <u>Data are representative of three independent experiments.</u> **(F, G)** In vivo cisplatin sensitivity studies were performed comparing the NT control group with the CD55-silenced group, and the graph shows the growth rate of tumors compared to the first day of cisplatin treatment. **(H)** <u>Hematoxylin/eosin stained slides of tumors excised from mice treated with cisplatin and vehicle controls.</u> *p< 0.5, **p< 0.01, ***p< 0.001. <u>Scale = 50 μm (30 μm in insets)</u>

Cisplatin resistance is a hallmark of endometrioid CSCs (Wiechert A, 2016), and based on the high expression of CD55 in CSCs and cisplatin resistant parental cells, we investigated whether CD55 inhibition impacts cisplatin resistance. CD55-silenced CSCs from cisplatin-naive cells lines (A2780, TOV112D), and PDX cells (EEC-4) had significantly higher sensitivity to cisplatin and lower survival rates at cisplatin doses from 2.5 to 50 μM, as compared to non-targeted control cells **(Fig. 2E; Supplementary Fig. S3A)**. Further, CD55-silenced CSCs demonstrated higher caspase 3/7 activity compared to non-targeted CSCs upon cisplatin treatment (2.5- 10 μM), indicating increased susceptibility to cisplatin-induced cell death **<u>(Supplementary Fig. S3B)</u>**. Similarly, CD55 inhibition led to increased sensitivity to cisplatin in cisplatin-resistant CP70 and HEC1a cell lines **(Supplementary Fig. S3C-D)**. To further validate the effect of CD55 silencing on cisplatin resistance, we injected CD55-silenced and control CSCs into a total of 45 mice at a concentration of 2 million cells/mouse, and waited until each mouse developed a 1 cm tumor **(Fig. 2F)**. As tumors reached the target size of 1 cm, mice were randomized 2:1 to receive cisplatin (2.5 mg/kg three times a week) and vehicle (DMSO) treatments, respectively. In vehicle control groups, mice with CD55-silenced tumors had significantly lower growth rates as compared to non-targeted controls **(Fig. 2F, G; Supplementary Fig. S3E)**. After 17 days of cisplatin treatment, tumors originating from CD55-silenced CSCs were more sensitive to cisplatin as compared to tumors originating from non-targeted CSC controls **(Fig. 2G; Supplementary Fig. S3E)**. Moreover, CD55-silenced tumors demonstrated higher degrees of cell death and tumor regression, inflammatory cell infiltrate, and fibrosis, as compared to non-targeted controls treated with cisplatin **<u>(Fig. 2H)</u>**. While CD59 expression was also increased in endometriod CSCs and cisplatin resistant cells, we did not observe any attenuation in CSC marker expression, self-renewal, or enhanced sensitivity to cisplatin upon shRNA silencing CD59 expression (**Supplementary Fig. S2E-F, S3F)**. These findings demonstrate that CD55 is necessary for the maintenance of cisplatin resistance in endometrioid CSCs and cisplatin resistant cells.

### CD55 is sufficient to drive CSC maintenance and cisplatin resistance

Based on the necessary role of CD55 in maintenance of self-renewal and cisplatin resistance, we investigated whether CD55 was sufficient to induce stemness and cisplatin resistance in non-CSCs and cisplatin-naive cells, both of which express low levels of CD55. We successfully transduced CD55 into non-CSCs of cisplatin naive cells (A2780, TOV112D) (Fig. 3A). <u>Upon CD55 overexpression, we observed an increase in protein expression of core pluripotency genes (**Fig. 3A**), We also observed an increase in NANOG and SOX2 mRNA levels upon CD55 overexpression **(Fig. 3B)**. Moreover, CD55 overexpression in non-CSCs led to significantly higher self-renewal and stem cell frequencies as compared to non-CSCs transduced</u> with empty vector (increased from empty vector to CD55 overexpression conditions as 1 in 33.8 to 1 in 18.8 for A2780 non-CSCs [p<0.05]; 1 in 23.9 to 1 in 12 for TOV112D non-CSCs [p<0.01]) **(Fig. 3C)**. Utilizing our NANOG promoter GFP reporter system, which allows for direct visualization of stemness, we demonstrated an increase in GFP signal upon CD55 overexpression **(Fig. 3D)**. Additionally, tumorspheres originating from CD55 overexpressing non-CSCs demonstrated a heterogeneous distribution of GFP signal, as compared to empty vector-transduced non-CSCs, which exhibited <u>low</u> GFP signal **(Fig. 3E)**. We further investigated whether CD55 overexpression was sufficient to induce cisplatin resistance. CD55 overexpressing non-CSCs had significantly higher rates of survival and lower levels of caspase 3/7 activity upon cisplatin treatment, as compared to non-CSCs with empty vector transduction **(Fig. 3F-G)**. These data demonstrate that CD55 is sufficient to induce CSC marker expression, self-renewal, and cisplatin resistance in non-CSCs.

**Figure 3.**
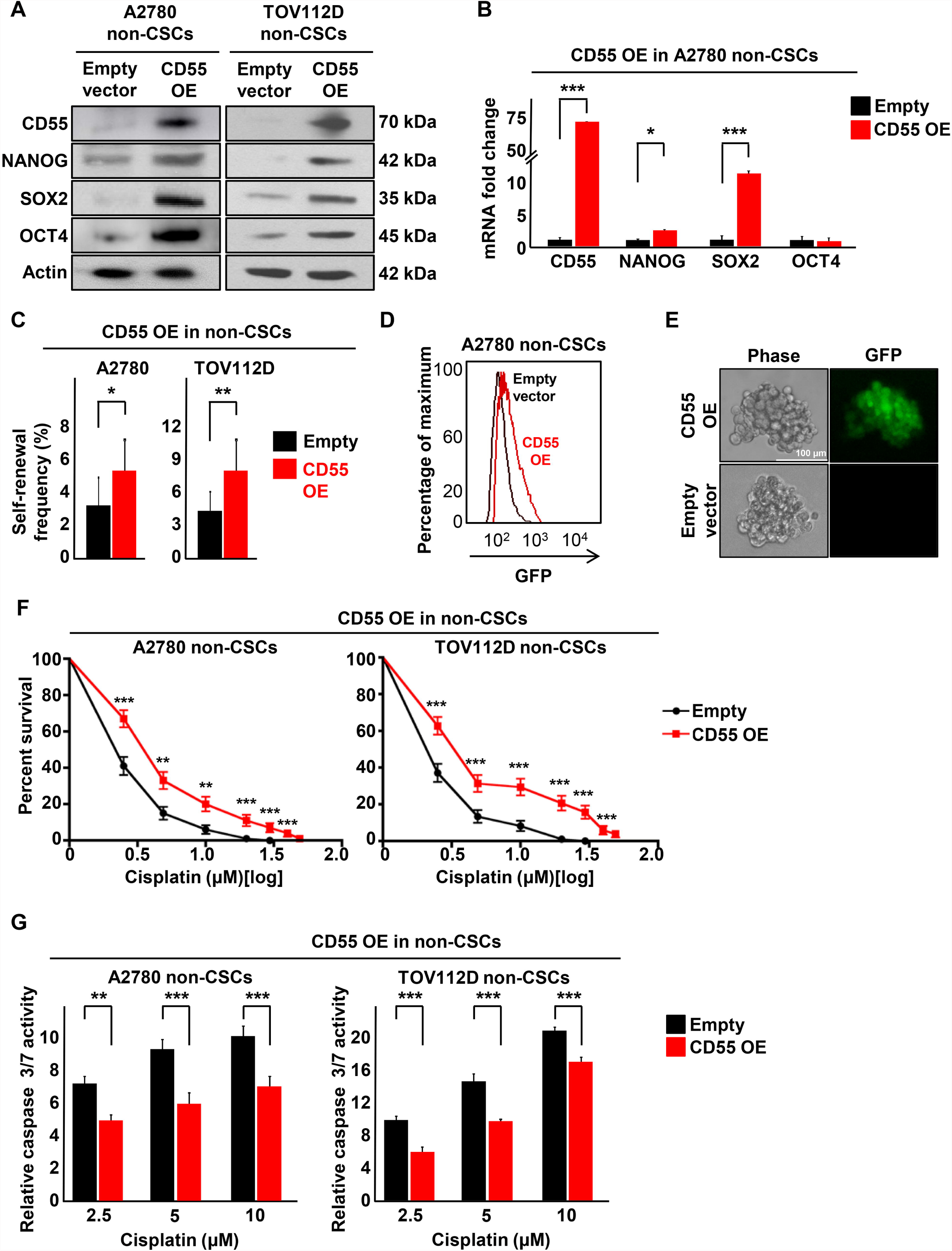
CD55 is sufficient to drive self-renewal and cisplatin-resistance in endometrioid non-CSCs. **(A)** Immunoblots of cisplatin-naïve non-CSCs with CD55 overexpression and empty vector controls were probed with CD55, NANOG, SOX2, and OCT4. Actin was used as loading control. <u>Data are representative of two independent experiments.</u> **(B)** mRNA expression was determined by qPCR and compared between CD55- overexpressing A2780 non-CSCs and empty vector control non-CSCs. Actin was used as a control. <u>Three technical replicates were used.</u> **(C)** Limiting dilution analysis plots of empty vector control compared with CD55 overexpression in cisplatin-naïve non-CSCs. The graph compares the estimates of the percentage of self-renewal frequency in these sorted populations with the corresponding p-values. <u>Data are representative of three independent experiments.</u> **(D)** A2780 non-CSCs transduced with CD55 overexpression and empty vector controls were flowed for GFP signal intensity, which indicates *NANOG* promoter activity. **(E)** Tumorsphere pictures for A2780 non-CSCs transduced with CD55 overexpression and empty vector controls. **(F)** CD55 overexpressing cisplatin-naïve non-CSCs and their empty vector controls were treated with 0-50 μM cisplatin and percent surviving cells were graphed. <u>Data are representative of three independent experiments.</u> **(G)** Relative caspase 3/7 activity for CD55 overexpressing cisplatin-naïve cells and their empty vector controls after treatment with cisplatin. <u>Relative caspase activities in cisplatin treated groups were calculated after normalizing the corrected readings to untreated controls in each group. Data are representative of two independent experiments and three technical replicates were used in each.</u> *p< 0.5, **p< 0.01, ***p< 0.001. <u>Scale bar: 100 μm.</u>

### CD55 regulates self-renewal and cisplatin resistance via a complement-independent mechanism

To interrogate the mechanism by which CD55 regulates these phenotypes, we first studied its canonical function, which is the inhibition of complement cascade. Since our cell culture conditions and NSG mice did not have complement proteins, we assessed this function by conventional BCECF-based cytotoxicity assay after incubating cells with normal human serum (NHS). We found that non-CSCs and cisplatin-naïve (A2780) cells, which had lower levels of CD55, had significantly higher amounts of BCECF leakage, as compared to their CSC and cisplatin-resistant (CP70) counterparts, respectively **(Supplementary Fig. S4A)**. Additionally, CD55+ A2780 cells demonstrated higher proliferative capacity at lower NHS doses, as compared to CD55- cells **(Supplementary Fig. S4B)**. However, complement treatment did not impact self-renewal or cisplatin resistance in CD55+ and CD55- cell populations **(Supplementary Fig. S4C-D)**. These data suggested that even though CD55+ cells are more resistant to complement-mediated cytotoxicity, addition of complement does not alter self-renewal or cisplatin resistance, which are regulated by complement-independent mechanisms.

### CD55 function depends on GPI-anchorage to lipid rafts

It has been reported that GPI-anchored proteins, including CD55, are localized to lipid rafts and can activate non-receptor tyrosine kinases (Shenoy-Scaria AM, 1992). First, we confirmed that CD55 localized to lipid rafts by coimmunolocalization with cholera toxin-B, a marker of lipid rafts **(Supplementary Fig. S4E)**. We investigated whether a GPI-deficient transmembrane CD55 (TM-CD55) construct can activate this signaling. We transduced non-CSCs with empty control, CD55 overexpression and TM-CD55 vectors, with the latter being a chimeric protein containing the extracellular portion of CD55 (amino acids 1-304) fused to the transmembrane and cytoplasmic domains of CD46 (amino acids 270-350) (Shenoy-Scaria AM, 1992). In non-CSCs transduced with CD55, the protein localized mainly to the lipid rafts, however TM-CD55 construct was distributed more uniformly on the membrane, with a significantly lower level of co-localization with the lipid raft marker (67.5% in CD55-transduced non-CSCs vs 18.7% in TM-CD55-transduced non-CSCs, p< 0.001) **(Fig. 4A-B)**. Despite the decreased lipid raft localization, non-CSCs transduced with TM-CD55 were resistant to complement-mediated cytotoxicity to the level of CD55-overexpressing non-CSCs **(Fig. 4C)**. However, TMCD55-transduced non-CSCs demonstrated lower self-renewal, stem cell frequencies (1 in 29.2 in empty vector-transduced, 1 in 11.8 in CD55-transduced [p< 0.001], 1 in 26.4 in TM-CD55-transduced [p< 0.01] non-CSCs), and cisplatin resistance, as compared to non-CSCs with CD55 overexpression **(Fig. 4D-E)**. Moreover, upon phosphatidylinositol-specific phospholipase C (PIPLC)-mediated cleavage of CD55 from membrane, CSCs became more sensitive to cisplatin **(Supplementary Fig. S4F)**. Collectively, these findings indicate that CD55 function depends on its anchorage to lipid rafts via the GPI-link.

**Figure 4.**
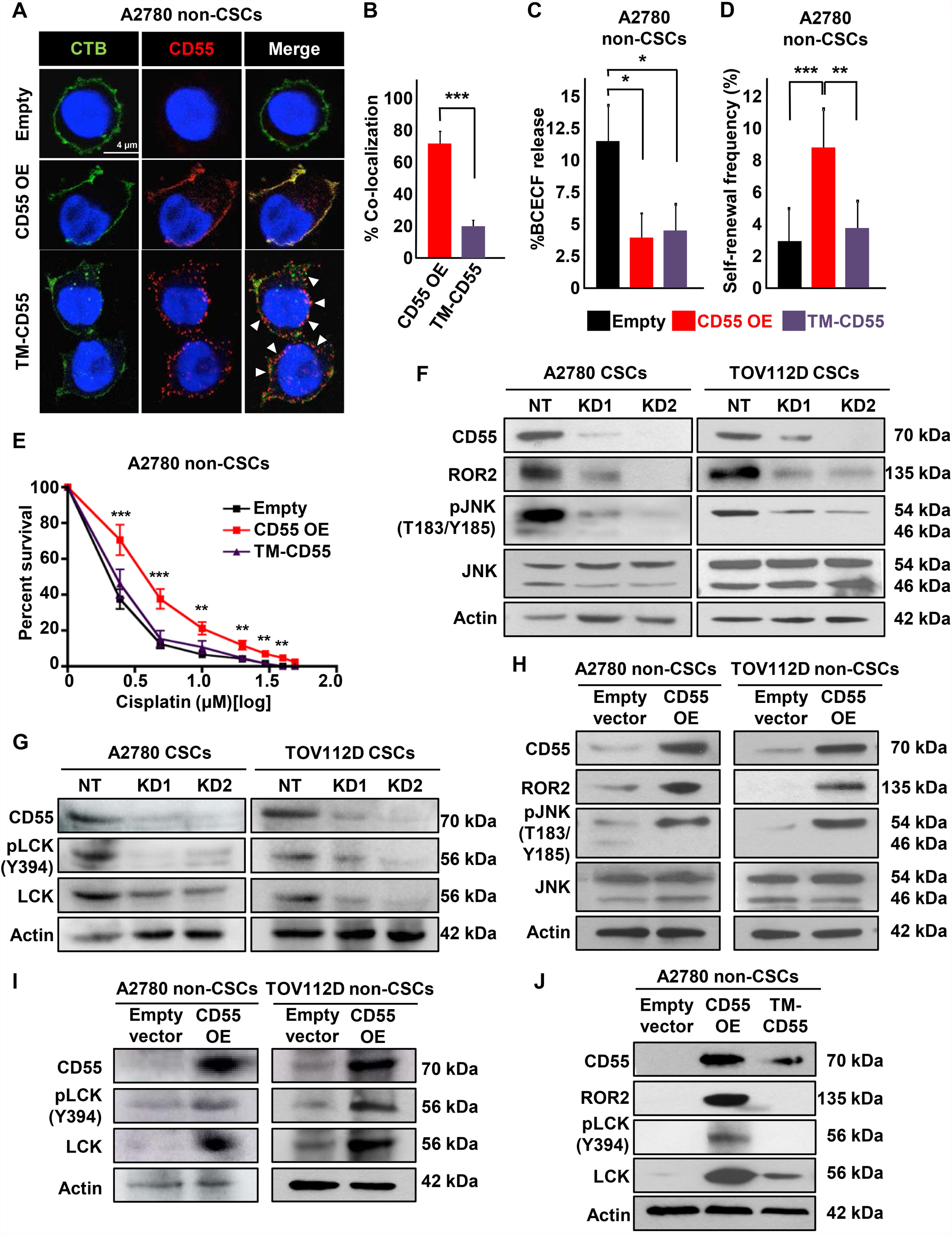
CD55 localization to lipid rafts is essential for its signaling via ROR2-JNK and LCK pathways. **(A)** Immunofluorescent staining of cisplatin-naïve non-CSCs transduced with CD55, GPI-deficient transmembrane (TM)-CD55, and empty vector control. The arrows point to areas where CD55 is not localized to lipid rafts. **(B)** The graph shows the percentage of CD55-cholera toxin B co-localization. <u>Data are representative of two independent experiments, quantifying >40 cells per group.</u> **(C)** Complement-mediated cytotoxicity as assessed by %BCECF dye release in A2780 non-CSCs transduced with CD55, TM-CD55, and empty vector control. <u>Data are representative of two independent experiments and three technical replicates were used.</u> **(D)** Limiting dilution analysis plots of CD55 empty vector control compared with CD55 overexpression and TM-CD55 constructs in cisplatin-naïve non-CSCs. **(E)** CD55 overexpressing cisplatin-naïve non-CSCs and their empty vector controls were treated with 0-50 uM cisplatin and percent surviving cells were graphed. <u>Data are representative of three independent experiments.</u> **(F, G)** Immunoblots of cisplatin-naïve CSCs silenced for CD55 using two shRNA constructs and a non-targeting control were probed with CD55, ROR2, pJNK (T183/Y185), JNK, pLCK (Y394), and LCK. Actin was used as a loading control. <u>Data are representative of two independent experiments.</u> **(H, I)** Cell lysates from cisplatin-naïve non-CSCs transduced with CD55 and empty vector control were probed for CD55, ROR2, pJNK (T183/Y185), JNK, pLCK (Y394), and LCK. Actin was used as a loading control. <u>Data are representative of two independent experiments.</u> **(J)** Immunoblots of cisplatin-naïve non-CSCs transduced with CD55, TM-CD55, and empty vector control were probed with CD55, ROR2, pLCK (Y394), and LCK. Actin was used as a loading control. *p< 0.5, **p< 0.01, ***p< 0.001. <u>Scale bar: 4 μm.</u>

### CD55 activates ROR2 and LCK kinases

To identify intracellular CD55 signaling pathways, we performed a receptor tyrosine kinase activation study using an antibody array against 71 tyrosine kinases **(Supplementary Fig. S4G)**. This screen revealed a decrease in levels of ROR2 and LCK in CD55-silenced A2780 CSCs, as compared to non-targeted CSC control **(Supplementary Fig. S4G)**. These results were further validated in cisplatin naive (A2780 and TOV112D) CSCs, in which CD55 inhibition led to decreased ROR2 and its downstream signaling via JNK pathway activation **(Fig. 4F)**. Additionally, CD55-silenced CSCs had lower levels of LCK and autophosphorylated active pLCK (Y394), as compared to non-targeted CSC controls **(Fig. 4G)**. CD55+ cells demonstrated higher activity of ROR2 and LCK pathways as compared to their CD55- counterparts **(Supplementary Fig. S4H)**. We could also induce the activation of these pathways with CD55 overexpression in non-CSCs **(Fig. 4H-I)**. While non-CSCs transduced with CD55 demonstrated active ROR2 and LCK signaling, these pathways were not induced in non-CSCs with TM-CD55 **(Fig. 3J)**. These data demonstrate that CD55 signals through ROR2 and LCK pathways and this signaling depends on its localization to lipid rafts in endometrioid tumors.

### LIME mediates intracellular CD55 signaling

As CD55 is an extrinsic protein tethered to the outer membrane via a GPI anchor, we searched for a transmembrane adaptor linking CD55 to signaling molecules located on the inner side of the membrane. We focused on known lipid raft adaptor proteins that were shown to interact with LCK. LIME (LCK interacting transmembrane adaptor) and PAG (protein associated with glycosphingolipid-enriched microdomains) emerged as candidates (Horejsí V, 2004; Ventimiglia LN, 2013). To identify CD55 interacting proteins, we performed an immunoprecipitation on endometrioid CSC lysates using CD55 antibodies and immunoblotted for LIME and PAG. We detected LIME but not PAG in the IP lysates **(Fig. 5A)**. To investigate the functional role of LIME in CD55 signaling, we silenced LIME in CSCs and found a decrease in the levels of ROR2, pLCK (Y394), and LCK **(Fig. 5B)**. Moreover, in LIME-silenced CSCs, CD55 was no longer able to interact with ROR2 and LCK to propagate the signaling **(Fig. 5C)**. We further assessed the impact of LIME inhibition on self-renewal and cisplatin resistance in CSCs. CSCs with LIME knockdown had lower levels of CSC markers, self-renewal, and stem cell frequencies (1 in 5.2 to 1 in 17.6 and 1 in 22.9, p<0.001), and higher sensitivity to cisplatin as compared to non-targeted control CSCs **(Fig. 5D-F)**. These data demonstrate that the transmembrane adaptor protein LIME is necessary for intracellular CD55 signaling and maintenance of self-renewal and cisplatin resistance.

**Figure 5.**
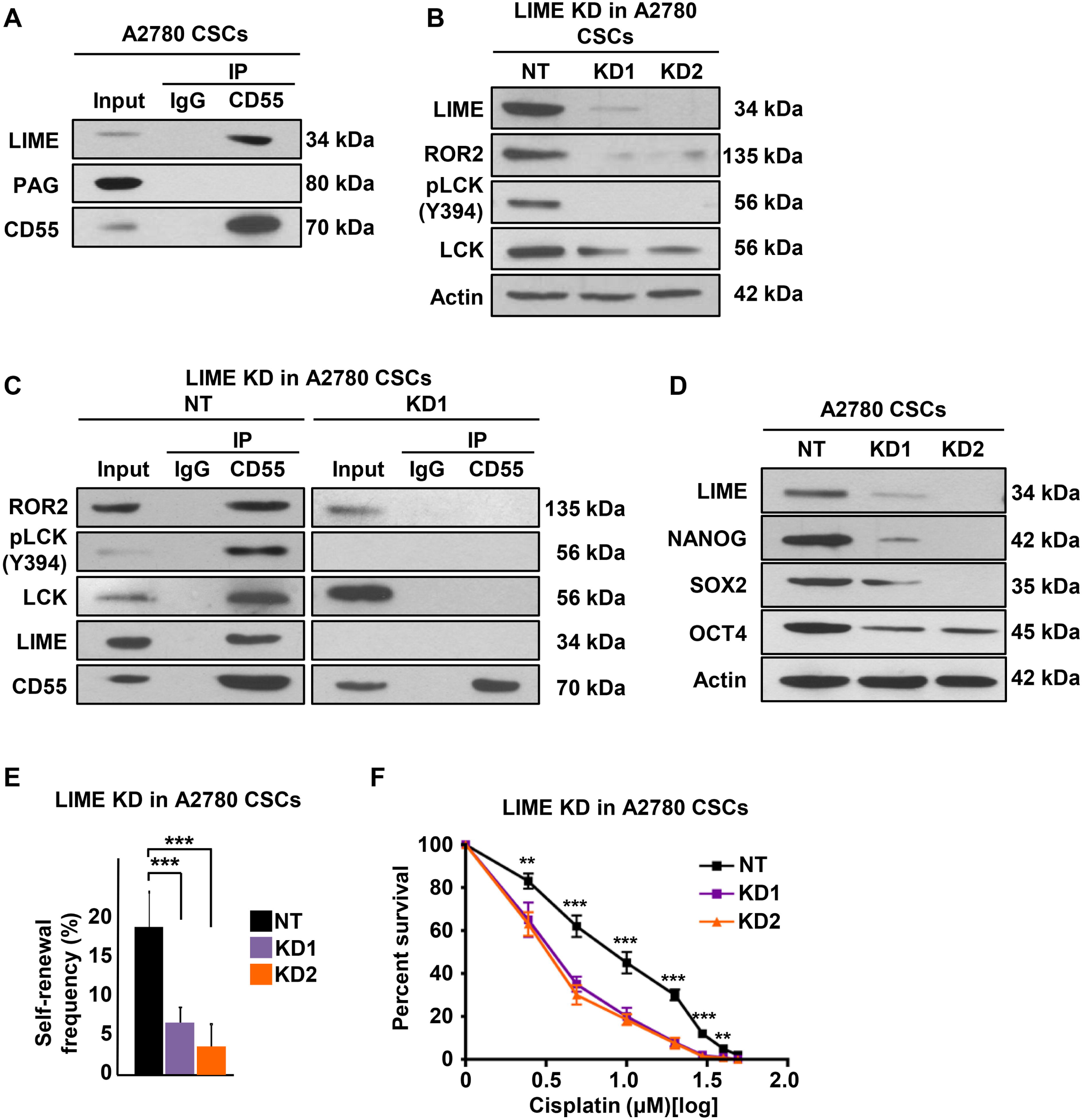
LIME is necessary for intracellular CD55 signaling. **(A)** Pull-down experiments with CD55 antibody were performed in cisplatin-naïve CSCs and elutes were probed for lipid raft adaptor proteins LIME and PAG. **(B)** Cell lysates from LIME silenced A2780 CSCs and their non-targeted (NT) controls were immunoblotted and probed with LIME, ROR2, pLCK (Y394), and LCK. Actin was used as loading control. **(C)** Pull-down experiments with CD55 antibody were performed in LIME-silenced and NT control cisplatin-naïve CSCs and elutes were probed for ROR2, pLCK (Y394), LCK, LIME, and CD55. **(D)** Immunoblots of cisplatinnaïve CSCs with LIME silencing and NT controls were probed with LIME, NANOG, SOX2, and OCT4. Actin was used as a loading control. **(E)** Limiting dilution analysis plots of LIME NT control compared with LIME sh1 and sh2 silencing constructs in cisplatin-naïve CSCs. **(F)** LIME silenced cisplatin-naïve CSCs and their NT controls were treated with 0-50 uM cisplatin and percent surviving cells are graphed. <u>All data are representative of two to three independent experiments.</u>

### CD55 activates ROR2-JNK signaling to maintain self-renewal

To elucidate the function of downstream CD55 signaling molecules, we first compared the expression of ROR2 between CSCs and non-CSCs **(Fig. 6A)**. CSCs of cisplatin-naïve cells (A2780 and TOV112D) had higher levels of ROR2 as compared to non-CSCs **(Fig. 6A)**. Both CSCs and non-CSCs demonstrated expression of p46 and p54 JNK isoforms, and the former was higher on CSCs. These isoforms were reported to be protein kinases with no functional difference (i.e. generated with differential mRNA processing) and suggested to be involved in ROR2 signaling (Oishi et al., 2003). In endometrioid tumor cells, only p54 JNK was phosphorylated and phospho-p54 JNK was higher on CSCs as compared to non-CSCs **(Fig. 6A)**. Based on our observation that CD55 knockdown decreased ROR2 levels and JNK pathway activity in CSCs, while CD55 overexpression induced ROR2-JNK signaling pathway, we assessed whether there was a direct or indirect link between CD55 and ROR2. We immunoprecipitated CD55 in A2780 and PDX (EEC-4) CSCs, and determined by immunoblotting that ROR2 was co-precipitated **(Fig. 6B)**. To investigate ROR2 signaling independently, we silenced ROR2 in CSCs, which in turn led to inhibition of p54 JNK phosphorylation, and decrease in levels of core pluripotency transcription factors (NANOG, SOX2, OCT4) **(Fig. 6C)**. We also showed a decrease in GFP intensity of CSCs, which indicated decreased NANOG promoter activity **(Fig. 6D)**. ROR2-silenced CSCs had significantly lower self-renewal and stem cell frequencies, as compared to non-targeted CSC controls (decreased from 1 in 4.4 to 1 in 20.7 and 1 in 31.7, p< 0.001) **(Fig. 6E)**. However, ROR2 inhibition did not impact cisplatin resistance in CSCs **(Fig. 6F)**. Collectively, these data indicate that CD55 interacts with transmembrane ROR2 protein and activates JNK pathway to maintain self-renewal.

**Figure 6.**
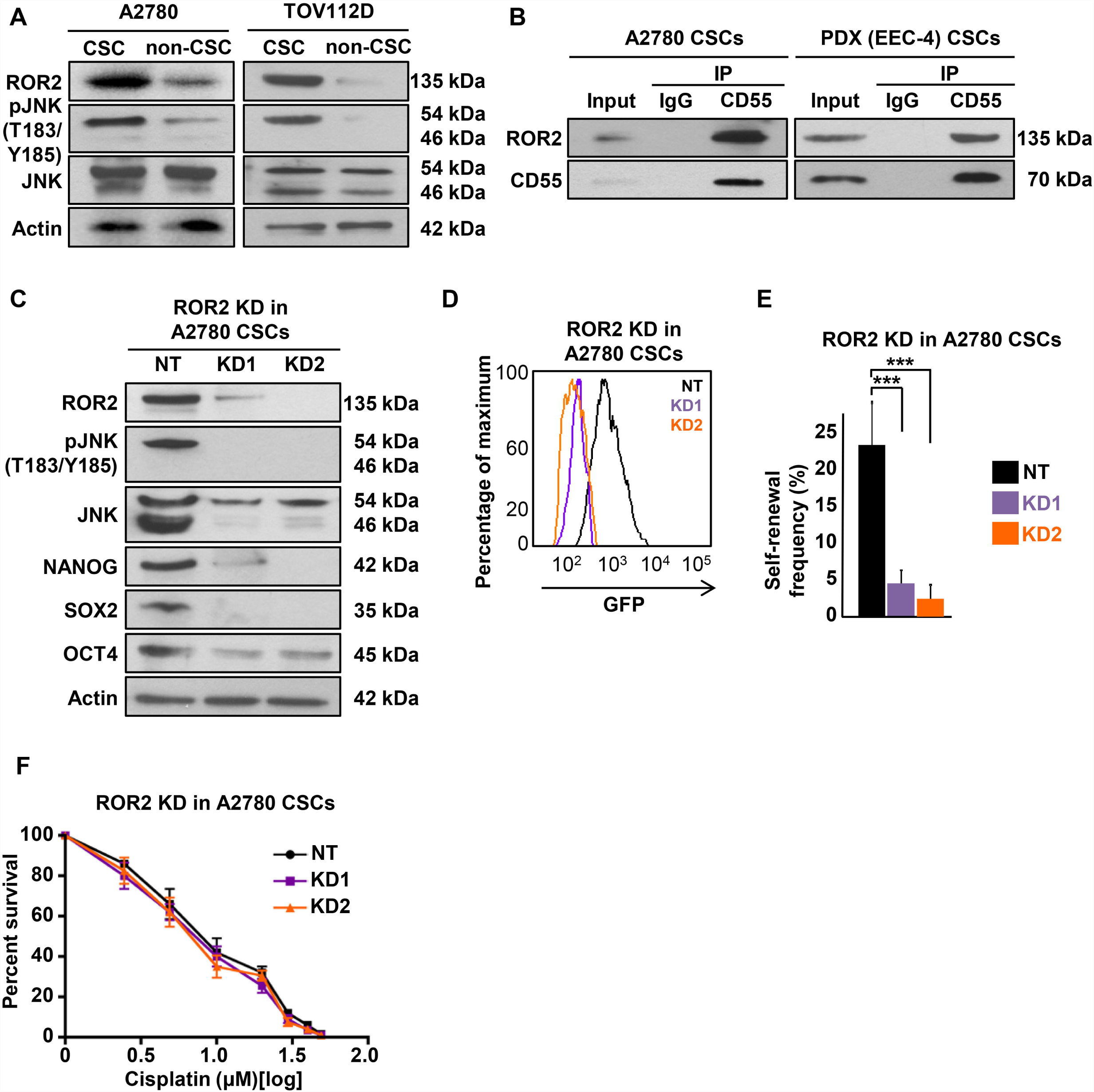
CD55 signals via ROR2-JNK pathway to regulate self-renewal. **(A)** Cell lysates from cisplatin naïve CSCs and non-CSCs were immunoblotted and probed for ROR2, pJNK (T183/Y185), and JNK. Actin was used as a loading control. **(B)** Pull-down experiments with CD55 antibody were performed in cisplatinnaïve CSCs and elutes were probed for ROR2. **(C)** Immunoblots of cisplatin-naïve CSCs silenced for ROR2 using two shRNA constructs and a non-targeting control were probed with ROR2, pJNK (T183/Y185), JNK, NANOG, SOX2, and OCT4. Actin was used as a loading control. **(D)** A2780 CSCs silenced for ROR2 and NT controls were flowed for GFP signal intensity, which indicates *NANOG* promoter activity. **(E)** Limiting dilution analysis plots of CD55 NT control compared with CD55 sh1 and sh2 silencing constructs in cisplatin-naïve CSCs. **(F)** ROR2 silenced cisplatin-naïve CSCs and their NT controls were treated with 0-50 μM cisplatin and percent surviving cells are graphed. <u>All data are representative of two to three independent experiments.</u> ***p< 0.001

### CD55 induces LCK signaling to drive cisplatin resistance

Based on the finding that CD55 signaling through ROR2-JNK pathway regulates self-renewal alone, we explored the role of LCK, which was the other kinase downregulated or induced with CD55 silencing and overexpression, respectively. CSCs and cisplatin resistant cells had higher levels of both pLCK (Y394) and LCK, as compared to their non-CSC and cisplatin-naïve counterparts, respectively **(Fig. 7A; Supplementary Fig. S5A)**. We did not detect phosphorylation of LCK at Y505 residue, which leads to inhibition of LCK, in any of these cells (data not shown). Moreover, when CD55 was immunoprecipitated in A2780 and PDX (EEC-4) CSCs, and cisplatin-resistant (CP70) cells, LCK and pLCK (Y394) were co-precipitated **(Fig. 7B; Supplementary Fig. S5B)**. To study the effects of LCK inhibition, we treated CSCs with a FYN/LCK inhibitor, saracatinib, and assessed self-renewal and cisplatin resistance. At 500 nM and 1 μM concentrations of saracatinib, we did not observe a significant change in self-renewal and stem cell frequencies (1 in 1.4 in DMSO control to 1 in 1.8 with 500 nM, and 1 in 2.6 with 1 μM saracatinib, p> 0.05) **(Supplementary Fig. S5C)**. However, CSCs treated with 1 μM saracatinib demonstrated significantly higher sensitivity to cisplatin, as compared to CSCs treated with cisplatin and DMSO **(Fig. 7C)**. In addition to pharmacologic inhibition, we silenced LCK in CSCs with three non-overlapping shRNA constructs **(Supplementary Fig. S5D)**. Similarly, LCK-silenced CSCs demonstrated significantly higher sensitivity to cisplatin **(Supplementary Fig. S5E)**. To investigate whether LCK is sufficient to drive these phenotypes, we transduced non-CSCs with LCK overexpression and empty control vectors. While LCK overexpression did not affect the levels of CSC markers and self-renewal in non-CSCs (stem cell frequencies: 1 in 24.1 in empty vector, 1 in 25.5 in LCK overexpression, p>0.05) **(Supplementary Fig. S5F-G)**, LCK overexpressing non-CSCs had significantly higher survival rates and lower caspase 3/7 activity levels as compared to non-CSCs with empty vector transduction **(Fig. 7D)**. To assess whether LCK inhibition can overcome CD55-induced cisplatin resistance, we treated CD55 overexpressing and empty vector-transduced non-CSCs with cisplatin and/or 1 μM saracatinib. While CD55-transduced non-CSCs were more resistant to cisplatin and had lower levels of caspase 3/7 activity, co-treatment with 1 μM saracatinib could overcome the resistance conferred by CD55 **(Fig. 7E-F, Supplementary Fig. S5H)**. To elucidate the particular mechanism of cisplatin resistance activated by CD55- LCK signaling, we performed a targeted screening of 31 genes involved in various mechanisms of platinum resistance including drug efflux, inactivation, and DNA repair **(Fig. 7G)**. When non-CSCs transduced with CD55 or LCK, and CSCs with CD55 silencing and saracatinib treatment were compared with their respective controls (i.e. empty vector, non-targeted control, and DMSO treatment, respectively), genes involved in DNA repair, including *MLH1* and *BRCA1* were found to be modulated by these modifications **(Fig. 7G, Supplementary Fig. S5I)**. Upon inhibition of *MLH1* and *BRCA1*, CSCs showed increased sensitivity to cisplatin (**Supplementary Fig. S5J-K**). These data indicate that CD55 signals through LCK pathway to induce cisplatin resistance via activation of DNA repair genes, and inhibition of this pathway with saracatinib can sensitize cells to cisplatin.

**Figure 7.**
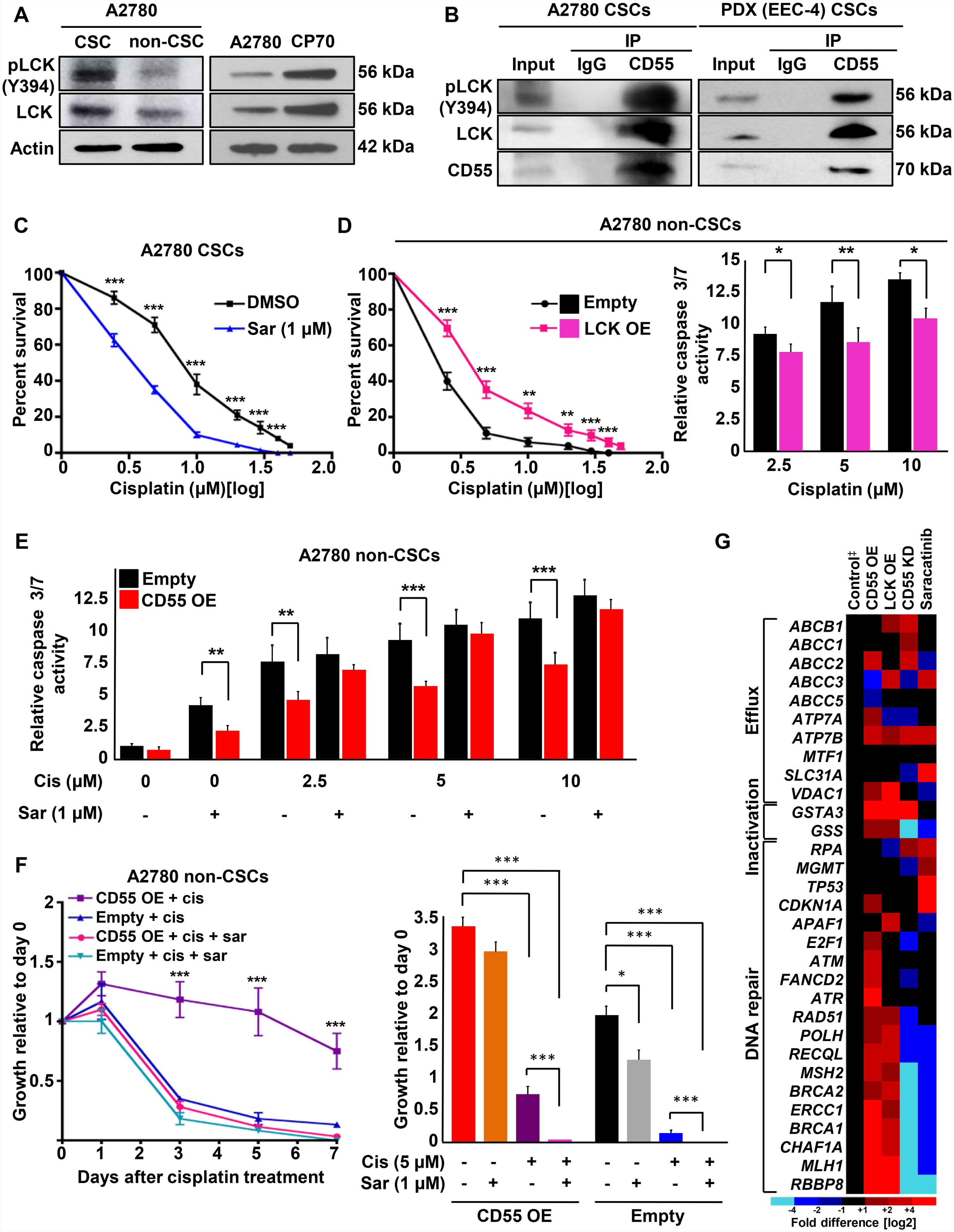
CD55 signals via LCK pathway to drive cisplatin resistance. **(A)** Cell lysates from cisplatin naïve CSCs and non-CSCs were immunoblotted and probed for pLCK (Y394) and LCK. Actin was used as a loading control. **(B)** Pull-down experiments with CD55 antibody were performed in cisplatin-naïve CSCs and elutes were probed for pLCK (Y394) and LCK. **(C)** Saracatinib (1 μM)-treated CSCs and their DMSO-treated controls were treated with 0-50 uM cisplatin and percent surviving cells are graphed. **(D)** LCK overexpressing cisplatinnaïve non-CSCs and their empty vector controls were treated with cisplatin and percent surviving cells and relative caspase 3/7 activity were graphed. **(E)** Relative caspase 3/7 activity for CD55 overexpressing non-CSCs and their empty vector controls treated with or without cisplatin (2.5-10 μM) and/or saracatinib (1 μM). **(F)** Growth curves for CD55-overexpressing non-CSCs and their empty vector controls treated with cisplatin with or without saracatinib. The graph shows growth relative to day 0. All data are representative of two to three independent experiments. **(G)** Targeted gene expression profiling of 31 genes involved in various mechanisms of cisplatin resistance was performed in cisplatin-naïve non-CSCs with CD55 or LCK overexpression, and CSCs with CD55-silencing or LCK inhibition with saracatinib.^‡^ ‡Emtpy vector control for non-CSCs, and non-targeted control for CSCs. <u>All data are representative of two independent experiments with three technical replicates.</u> *p< 0.5, **p< 0.01, ***p< 0.001

Collectively, these findings demonstrate that CD55 is GPI-anchored to lipid rafts, and signals via LIME to activate ROR2-JNK and LCK pathways to regulate self-renewal and cisplatin resistance, respectively **(Fig. 8)**.

**Figure 8.**
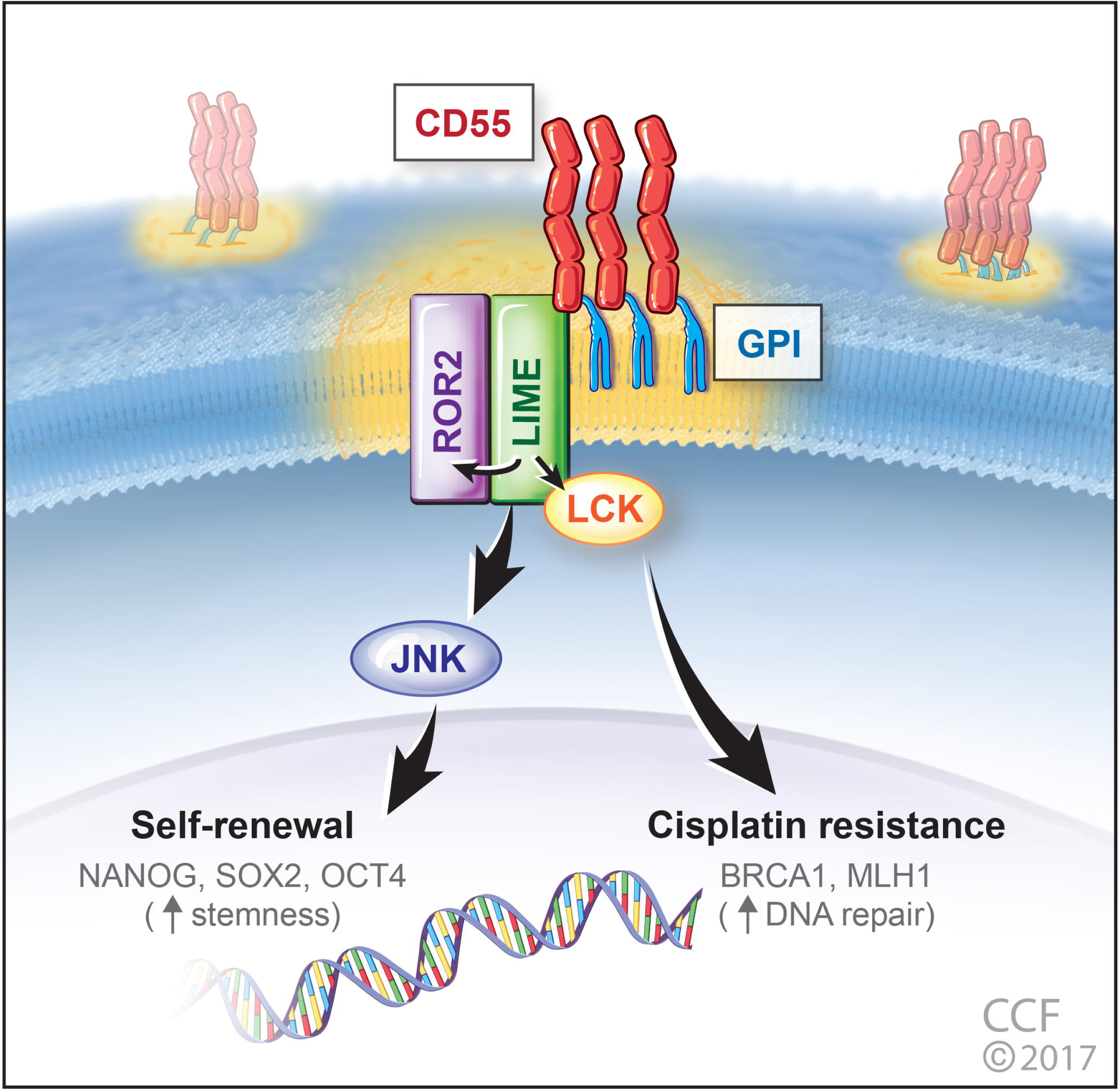
CD55 regulates self-renewal and cisplatin resistance in endometrioid tumors. CD55 is glycophosphatidylinositol (GPI)-anchored to lipid rafts and through LIME-mediated signaling, it activates ROR2-JNK pathway to regulate self-renewal, and LCK pathway to induce the expression of DNA repair genes and drive cisplatin resistance.

## DISCUSSION

These data provide the first evidence of CD55 signaling in a complement-independent manner in solid tumors to regulate self-renewal and therapeutic resistance. While previous efforts have identified CD55 as a prognostic marker in several cancers, our data provide mechanistic insight into a bifurcating signaling network that regulates self-renewal via ROR2/JNK signaling and cisplatin resistance via LCK signaling. <u>Our functional studies demonstrate that CD55 is necessary and sufficient for CSC maintenance. Mechanistically, CD55 regulates the protein expression of the key pluripotency transcription factors NANOG, SOX2, and OCT4, master regulator of CSC self-renewal. CD55 regulates NANOG and SOX2 protein expression at the transcriptional level (Qiu et al.,</u> 2010<u>), however regulation of OCT4 expression is more complex and indicative of post-transcriptional regulation, either at the level of protein synthesis or protein stability via phosphorylation, ubiquitination, and/or SUMOylation (Saxe et al., 2009; Yao et al., 2014).</u> Insights into CSC biology have uncovered a series of molecular mechanisms that individually regulate self-renewal and therapeutic resistance but few signaling networks have the capacity to impact both processes. CD55 represents one such signaling hub that both pathways originate from and hence represents an attractive therapeutic target in endometrioid cancers. In our pre-clinical studies, we observed that Saracatinib sensitized CSC to cisplatin and overcame CD55-induced chemoresistance but did not alter self-renewal. These studies suggest that the downstream CD55 signaling can be targeted with currently available agents but highlights the need for the development of CD55 inhibitors that attenuate both self-renewal and therapeutic resistance. While the mechanism that govern self-renewal and therapeutic resistance have traditionally posed barriers to effective treatment with conventional chemotherapy, CD55 intracellular signaling represents a central target which offers an opportunity to prevent recurrence and associated morbidity and mortality in patients with endometrioid cancer.

## MATERIALS AND METHODS

### Cell culture

The isogenic endometrioid ovarian cancer cell lines A2780 (cisplatin naive) and CP70 (cisplatin resistant) were cultured in log-growth phase in DMEM medium supplemented with 10% heat-inactivated fetal bovine serum (HI-FBS) at 37 °C in a humidified atmosphere (5% CO_2_). Endometrioid TOV112D ovarian cancer cell line was cultured in a 1:1 mixture of MCDB 105 medium and Medium 199, supplemented with 15% HI-FBS. Patient-derived primary endometrioid endometrial cancer xenograft (PDX) EEC-4 was a kind gift from Dr. Kim's laboratory and maintained in RPMI 1640 with 10% HI-FBS (Unno K, 2014). Cisplatin-resistant primary endometrial cancer cell line HEC1a was cultured in modified McCoy's 5a medium. Cell lines were obtained from American Type Culture Collection (ATCC) and authenticated by short tandem repeat (STR) DNA profiling analysis. At 70-90% confluence, trypsin (0.25%)/EDTA solution was used to detach cells for passaging and further experiments until passage number 15. Primary uterine endometrioid cancer cells, UTE-1 and UTE-2, were freshly dissociated from surgical specimens. Cisplatin was obtained from Cleveland Clinic Hospital pharmacy and 1 mg/mL stock solutions were stored at 4 °C. Saracatinib (AZD0530) was obtained from Selleck Chemicals and 50 uM stock solutions were stored at -20 °C.

### Flow cytometry and high-throughput flow screen

Endometrioid tumor cells at a concentration of 1 million cells/mL were sorted on BD FACS Aria II to isolate cancer stem cells (CSCs) and non-CSCs. For NANOG-GFP sorting, GFP high and low populations were sorted from NANOG-GFP promoter transduced stable A2780/CP70 cells as previously described (Wiechert A, 2016). The antibodies used for FACS analysis were: APC-conjugated integrin α6 (1:100, BD Biosciences), and APC-conjugated CD55 (1:100, BD Biosciences). Appropriate isotype controls were used to set gates. Data analysis was performed using the Flowjo software (Tree Star, Inc., Ashland, OR).

For the high-throughput flow cytometry screening, we used BD Lyoplate Human Cell Surface Marker Screening Panel which was purchased from BD Biosciences. The panel contains 242 purified monoclonal antibodies to cell surface markers and both mouse and rat isotype controls for assessing background signals. For screening procedure, A2780 and CP70 NANOG-GFP cells were prepared in single cell suspensions in BD Pharmingen Stain Buffer with the addition of 5 mM EDTA. The screening was performed as previously described (Thiagarajan PS, 2015). A2780 and CP70 NANOG-GFP cells were stained with DRAQ5 (eBioscience, San Diego, CA) and pacific blue dyes (Life Technologies Grand Island, NY), respectively. The cells were then pooled and plated in 96-well plates (BD Biosciences, Franklin Lakes, NJ). Reconstituted antibodies were added to the wells as per the human lyoplate screening panel. After the washes, cells were stained with APC-labeled goat anti-mouse IgG secondary antibody (BD Biosciences, Franklin Lakes, NJ) and stained with a live/dead fixable blue dead cell stain kit (Life Technologies, Grand Island, NY). Cells were analyzed on a Fortessa HTS system (BD Biosciences, Franklin Lakes, NJ). Data were analyzed with FlowJo software and appropriate isotype controls were used to detect positive immunoreactivity.

### Immunoblotting and immunoprecipitation

For immunoblots, whole cell protein extracts were obtained with lysis of cells in 20 mM Tris-HCl (pH 7.5), 150 mM NaCl, 1 mM Na_2_EDTA,1 % NP-40, 1 mM EGTA, 1% sodium pyrophosphate, 1 mM β-glycerophosphate, 1 mM sodium orthovanadate, 1 ug/mL leupeptin, 20 mM NaF and 1 mM PMSF. Protein concentrations were measured with Bradford reagent (BIO-RAD, CA). Proteins in lysates (30-50 ug of total protein) were resolved by 10% SDS-PAGE and transferred to nitrocellulose membrane. Membranes were incubated overnight at 4 °C with primary antibodies against CD55 (1:1000) (Santa Cruz, CA), CD59 (1:1000) (Abcam), CD46 (1:1000) (Santa Cruz, CA), NANOG (1:500) (Cell Signaling), SOX2 (1:500) (Cell Signaling), OCT4 (1:500) (Cell Signaling), ROR2 (1:1000) (BD Biosciences), pJNK (1:1000) (T183/Y185) (Cell Signaling), JNK (1:1000) (Cell Signaling), pLCK (Y394) (1:1000) (BD Biosciences), LCK (1:1000) (Santa Cruz, CA), LIME (1:1000) (Invitrogen), PAG (1:1000) (Genetex), and β-actin (1:1000) (Cell Signaling). Secondary anti-mouse or anti-rabbit IgG antibodies conjugated to horse radish peroxidase (HRP) (1:2000) (Thermo, Rockford, IL) were used and immunoreactive bands were visualized using the ECL plus from Pierce (Rockford, IL, USA).

For immunoprecipitation, cells were lysed in 0.5% Triton X-100, 50 mM Tris (pH 7.6), 300 mM NaCl, 1 mM sodium orthovanadate, 5 mM EDTA, 10 ug/mL leupeptin, 10 ug/mL aprotinin, 10 mM iodoacetamide, and 25 ug/mL p-nitrophenyl guanidinobenzoate as previously described (Shenoy-Scaria AM, 1992). The lysates were spun at 12,000xg for 15 min at 4 °C. Supernatants were incubated with rabbit anti-human CD55 primary antibody (SantaCruz, CA) and the corresponding antibody control for 1 hour at 4 °C. Protein A/G agarose beads (Santa Cruz, Dallas, TX) were added to lysates which were subsequently incubated on a rotating mixer overnight at 4 °C. The beads were then washed 3-4 times at 4 °C, and Laemmli sample buffer was added to the beads and boiled for 5 minutes. Immunoblotting was performed using the indicated primary antibodies described above.

### Quantitative real time PCR (qPCR)

Total RNA was extracted from cancer stem and non-stem cells, CD55 knockdown and overexpressing cells and their respective controls, saracatinib treated cells and LCK overexpressing cells using RNeasy kit (Quiagen). For mRNA analysis, cDNA was synthesized from 1 ug of total RNA using the Superscript III kit (Invitrogen, Grand Island, NY). SYBR Green-based real time PCR was subsequently performed in triplicate using SYBR-Green master mix (SA Biosciences) on Applied Biosystems StepOnePlus real time PCR machine (Thermo). For analysis, the threshold cycle (Ct) values for each gene were normalized to expression levels of β-actin. The primers used were:

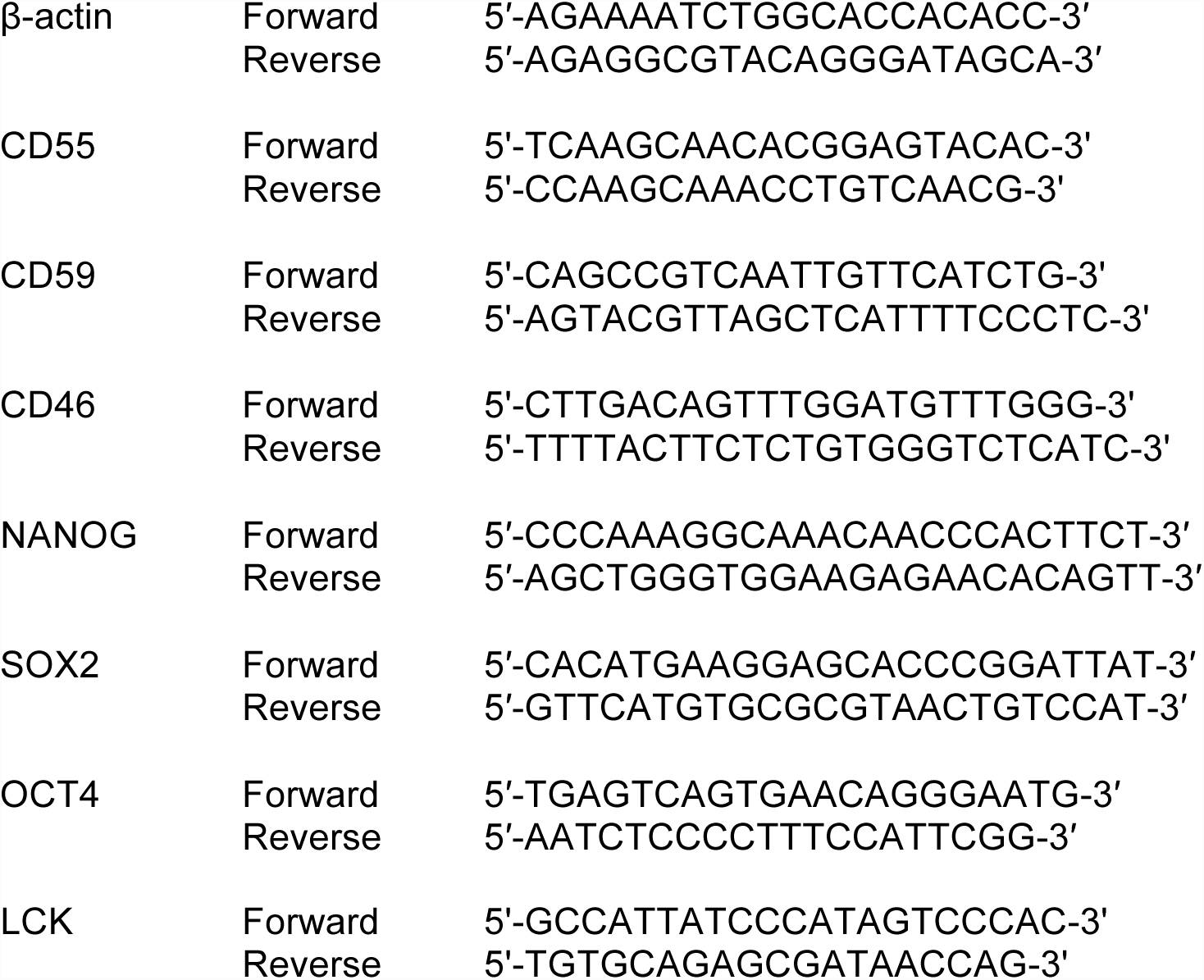

### Limiting dilution assays

For tumorsphere formation assays, BD FACS Aria II sorter was used to sort cells in duplicate rows of serial dilutions into 96-well ultra low attachment plates (Corning, Tewkesbury, MA, USA) with 200 uL serum-free DMEM/ F12 medium per well supplemented with 10 ng/mL epidermal growth factor (Biosource, Grand Island, NY, USA), 20 ng/mL basic fibroblast growth factor (Invitrogen), 2% B27 (Invitrogen), 10 ug/mL insulin, and 1 ug/mL hydrochloride (Sigma). Tumorspheres were counted in 2 weeks under a phase contrasted microscope and data was analyzed by Extreme Limited Dilution Analysis (ELDA) platform to determine stem cell frequency (http://bioinf.wehi.edu.au/ software/elda/) (Hu Y, 2009).

### Lentivirus production and infection

Lentiviral short hairpin RNAs (shRNAs), and CD55- and LCK-transducing lentiviruses were prepared as we previously reported (Lathia JD, 2010; Lathia JD, 2014). HEK 293T/17 cells were co-transfected with the packaging vectors pMD2.G and psPAX2 (Addgene, Cambridge, MA), and lentiviral vectors directing expression of shRNA specific to *CD55* (TRCN0000057167, TRCN0000057377), *CD59* (TRCN0000057108, TRCN0000057112), *ROR2* (TRCN0000001490, TRCN0000001491), *LIME* (TRCN0000257009, TRCN0000257011), LCK (TRCN0000001598, TRCN0000001599, TRCN0000001600), *MLH1* (TRCN0000040053, TRCN0000040056), *BRCA1* (TRCN0000039834, TRCN0000039835), a non-targeting (NT) control shRNA (SHC002), and overexpression vector for *CD55*, *LCK*, or an empty vector (Applied Biological Materials, Richmond, BC, Canada). Media of the HEK 293T/17 cells were changed 18 hours after transfection, and viral particles were harvested at 48 hours via concentration with polyethylene glycol precipitation, and stored at -80 °C for future use. Viral infections were performed in endometrioid tumor cell lines and PDX cells, and following transduction, cells were selected using 2-5 ug/mL puromycin.

### Cell survival and caspase 3/7 activity assays

Endometrioid CSCs, non-CSCs, and cisplatin resistant cells were plated in 12-well plates at 50,000 cells/well density and treated on the next day with cisplatin at the doses of 0-50 uM, and/or 1 uM saracatinib. The number of live cells in control and treatment groups were manually counted using hemocytometer at days 5 and 7 using Trypan blue dye exclusion as a live cell marker. Percentages of surviving cells at different treatment doses were normalized to the untreated control.

Apoptosis was measured using the Caspase-Glo 3/7 Assay kit (Promega, Southampton, UK) according to the manufacturer's instructions. Measured caspase activities were corrected for viable cell density as assessed by CellTiter-Glo (Promega, Southampton, UK). Relative caspase activities in cisplatin treated groups were calculated after normalizing the corrected readings to untreated controls in each group.

### Xenograft studies

NOD severe combined immunodeficient (SCID) IL2R gamma (NSG) mice were purchased from the Biological Response Unit (BRU) at the Cleveland Clinic and maintained in microisolator units with free access to water and food. For *in vivo* tumor initiation assay, CD55 knockdown and NT control A2780 CSCs were transplanted subcutaneously in serial dilutions of 1000, 10000, and 100000 cells (5 mice per group) into the right subcutaneous flank of female mice at 6 weeks of age. Mice were monitored every day until the endpoint of day 30, when the tumors that were palpable with a cross-sectional area >2 mm^2^ were taken as a positive read. Mice were euthanized and the tumors were resected. The stem cell frequencies were calculated using the ELDA algorithm as described above.

For the cisplatin treatment studies, NSG mice were injected subcutaneously with CD55 knockdown and NT control A2780 CSCs (15 mice per group). Each mouse was transplanted with 2 million cells to ensure tumor formation and tumors were allowed to grow to 1 cm in largest diameter. Then, mice were randomized into two groups, and one group (10 mice) was treated intraperitoneally with cisplatin (2.5 mg/kg, three times per week), while the other group (5 mice) received vehicle (DMSO). Tumor size was assessed at indicated time points by caliper measurements of length and width and the volume was calculated according to the formula (length x width^2^/2). Treatments were continued until day 14 in vehicle, and day 17 in cisplatin arms at which time the average tumor size reached 2 cm. Mice were euthanized and the tumors were resected for staining with hematoxylin/eosin. All mouse procedures were performed under adherence to protocols approved by the Institute Animal Care and Use Committee at the Lerner Research Institute, Cleveland Clinic.

### Complement-mediated cytotoxicity assay

A2780/CP70 parental cell, CSC, and non-CSC cytotoxicity after incubation with serum was assessed by BCECF (2′,7′-bis-(2-carboxyethyl)-5-(and-6)-carboxyfluorescein) leakage assay as previously described (Li Y, 2012). First, 2x10^5^ cells were labeled by incubation with 5 uM of BCECF-AM (Invitrogen) for 30 minutes at 37 °C. After washing, the labeled endometrioid tumor cells were incubated with 10-30% normal human serum (NHS) or respective controls in 100 uL of GVB^++^ buffer for another 30 minutes at 37 °C. Then, supernatants were collected, and BCECF dye release was measured by a fluorescence microtiter plate reader (Molecular Devices) with excitation and emission wavelengths of 485 nm and 538 nm, respectively. The percentage of BCECF release (indicative of complement mediated injury) was calculated with the following formula: [(AB)/(C-B)]x100%; where A represents the mean experimental BCECF release, B represents the mean spontaneous BCECF release (in the absence of serum), and C represents the mean maximum BCECF released that was induced by incubating cells with 0.5% Triton X.

### Immunocytochemistry

To visualize the expression and localization of CD55 and cholera toxin B, a lipid raft marker, A2780 and TOV112D CSCs were plated on coverslips placed in a 6-well plate. After 12-16 hours, the cells were fixed for 15 minutes with 4% paraformaldehyde at room temperature (RT), and washed three times with PBS. After washing, cells were incubated with A488-conjugated cholera toxin B (Invitrogen) for 15 minutes, and washed again for three times. Then, they were blocked in 5% goat serum with 1 mg/mL BSA for 2 hours. Mouse monoclonal CD55 antibody (Santa Cruz, CA) was used to stain cells overnight at 4 °C. The following day, cells were washed three times with PBS for 5 minutes and A647-conjugated goat anti-mouse secondary antibody (Invitrogen) was applied for 1 hour at RT. After secondary antibody incubation, cells were washed three times with PBS for 5 minutes each and counterstained with 4,6-diamidino-2-phenylindole (DAPI) for 5 minutes. Afterwards, cells were washed three times with PBS for 5 minutes each. The coverslips were mounted using 50% glycerol, and cells were imaged using <u>Leica TCS SP8 Confocal/Multi-Photon high speed upright microscope.</u>

### Generation of GPI-deficient transmembrane CD55 construct

A GPI-deficient transmembrane CD55 (TM-CD55) construct was generated as described elsewhere (Shenoy-Scaria AM, 1992). Briefly, TM-CD55 consisted of the extracellular portion of CD55 (amino acids 1-304) fused to the transmembrane and cytoplasmic domains of CD46 (membrane cofactor protein) (amino acids 270-350). First, the region of CD55 cDNA from amino acids 1 to 304 was amplified using the specific primers (forward: 5'-ATGACCGTCGCGCGGCC-3'; reverse: 5'-AACATTTACTGTGGTAGGTTTC-3'). Next, the region of CD46 cDNA from amino acids 270 to 350 was amplified using the specific primers with a stop codon added in the primer (forward: 5'-TGTGACAGTAACAGTACTTGG-3'; reverse: 5'-TCAAATCACAGCAATGACCC-3'). Then, the two PCR products were mixed in equal proportions and a single fusion/chimeric PCR product was generated using Mega PCR. The generated chimeric cDNA PCR product was cloned into pENTR/Directional TOPO vector and then recombined into pLenti-CMV-Puro-Dest vector (Addgene). For transformation, competent E. Coli strain DH5α was used to introduce 100 ng plasmid via heat shock at 42 °C for 45 seconds. Bacterial colonies resistant to ampicillin were selectively grown, and lentivirus was produced and cells were infected as described above.

### Phosphatidylinositol-specific phospholipase C (PIPLC) treatment

To release CD55 from the lipid rafts, CSCs were treated with the enzyme PIPLC (Sigma) at a final concentration of 4 U/mL, and compared with untreated cells. One unit of PIPLC liberates one unit of acetylcholinesterase per minute at pH 7.4 at 30 °C.

### Receptor tyrosine kinase array

For the receptor tyrosine kinase (RTK) activation study, a RayBio antibody array against 71 unique tyrosine kinases (Raybio AAH-PRTK-1-4) was used according to the manufacturer's protocol. Cell lysates (1 mg) from A2780 CSCs transduced with NT and two non-overlapping CD55 knockdown shRNAs were added to each membrane. Spot quantitation was done using Image J, and mean densities were calculated for each spot in a duplicate, and normalized to the densities of background and positive control dots.

### Gene expression profiling

To identify genes responsible for CD55-mediated regulation of cisplatin resistance, we performed a targeted screening of 31 genes involved in various mechanisms of platinum resistance including drug influx/efflux, inactivation, and DNA repair (Galluzzi L, 2014). RNA lysates from A2780 CSCs with CD55 knockdown vs NT control, saracatinib vs vehicle treatment, and non-CSCs with CD55 overexpression vs empty control, LCK overexpression vs empty control were used to perform serial RT-PCRs in triplicates and the relative amount of cDNA was calculated by the comparative CT method using actin sequence as the loading control. Fold-differences in gene expression were plotted in a heat-map. Primer sequences are listed below:

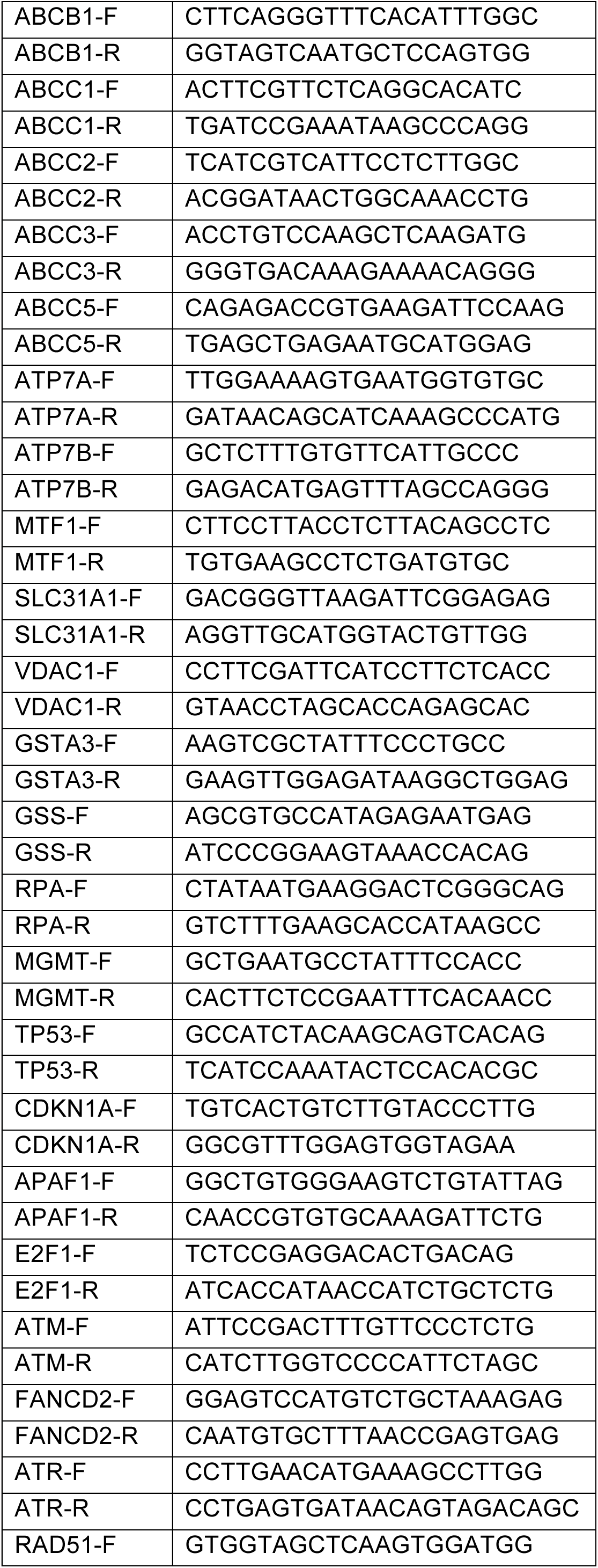
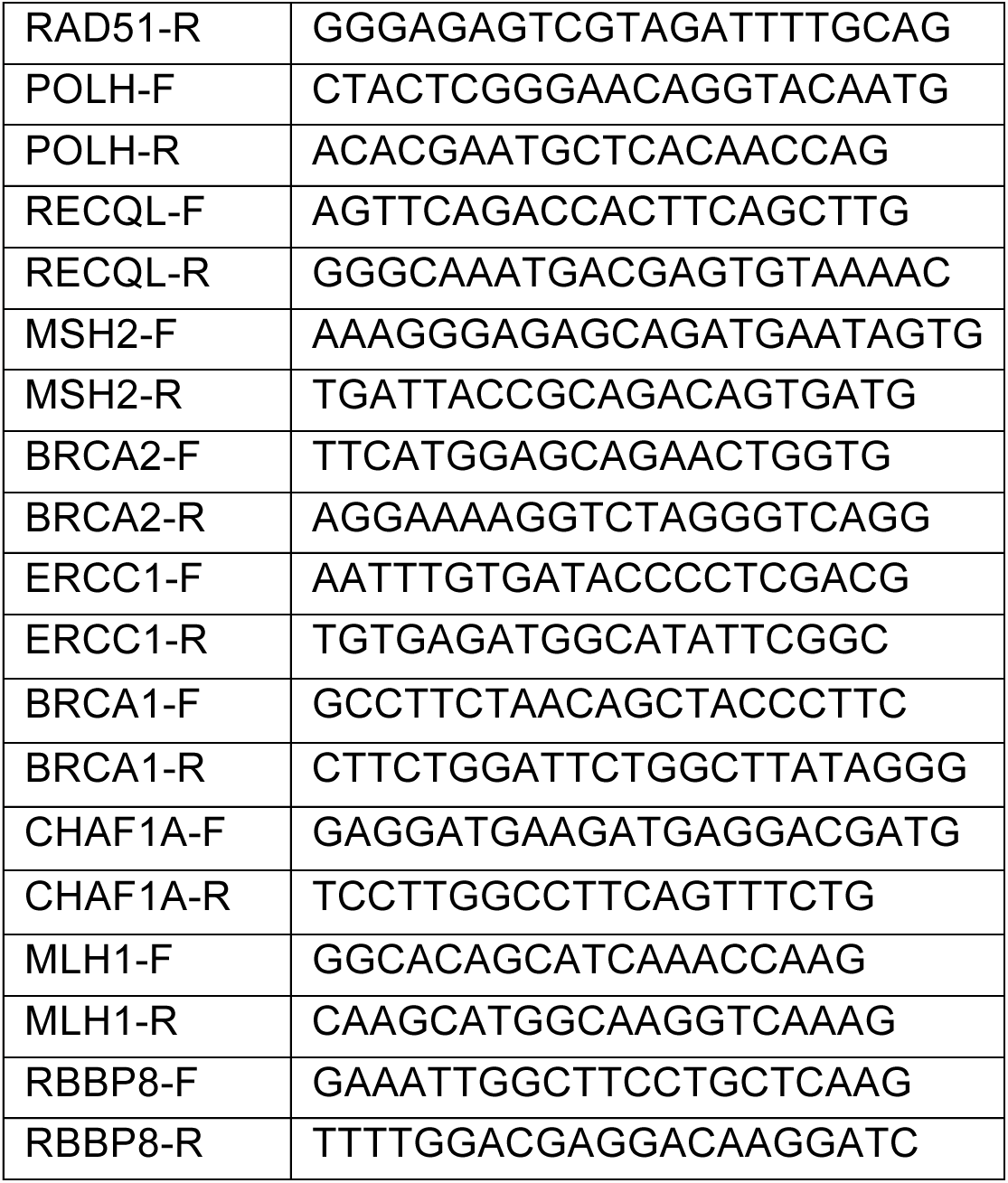

### Statistical Analysis

Values reported in the results are mean values +/- standard deviation. One-way ANOVA was used to calculate statistical significance, and the p-values are detailed in the text and figure legends.

## Author contributions

Conception and design (CS, OR, JDL), Financial support (OR, JDL), Collection and/or assembly of data (CS, AW, VSR, RA, EC, PST, JSH, YL, AC, AJ, DL, YP), Provision of reagents (ABN, JJK, AD, FAK, HM, FL, PGR, CM, KRM), Data analysis and interpretation (CS, VSR, RA, FL, OR, JDL), Manuscript writing (CS, OR, JDL), Final approval of manuscript (all authors).

## Acknowledgements

We thank Dr. Thomas McIntyre, and the members of the Reizes and Lathia laboratories for insightful discussion and constructive comments on the manuscript. We thank Cathy Shemo, Patrick Barrett, Sage O‘Bryant, Joseph Gerow, and Eric Schultz for flow cytometry assistance. We thank Amanda Mendelsohn from Cleveland Clinic Center for Medical Art and Photography for her exceptional help with the illustration of our model. This work was funded by National Institutes of Health grant CA191263 to O.R. and J.D.L. The work in the Reizes laboratory is also supported by the Cleveland Clinic Foundation, Clinical and Translational Science Collaborative of Cleveland (UL1TR000439), Case Comprehensive Cancer Center (P30 CA043703), the Cleveland Clinic VeloSano Bike Race, and the Laura J. Fogarty Endowed Chair for Uterine Cancer Research (O.R.). The Lathia laboratory also receives funding from the National Institutes of Health grants NS089641, NS083629, and CA157948; a Distinguished Scientist Award from the Sontag Foundation, a Research Scholar Award from the American Cancer Society, Blast GBM, the Cleveland Clinic VeloSano Bike Race, and the Case Comprehensive Cancer Center. This work utilized the Leica SP8 confocal microscope that was purchased from National Institutes of Health SIG grant 1S10OD019972-01.

<u>The authors declare no competing financial interests.</u>

## Supplementary Information

Supplementary Fig. S1 includes additional data on high CD55 expression in CSCs of cisplatin-naive and resistant cells, including freshly dissociated primary endometrioid uterine cancer specimens. Supplementary Fig. S2 and S3 show additional data supporting the role of CD55 in maintaining endometrioid tumor self-renewal and cisplatin resistance. Supplementary Fig. S4 provides data supporting the complement-independent function of CD55 and Supplementary Fig. S5 shows that CD55 induces DNA repair genes to regulate cisplatin resistance.

## FIGURE LEGENDS

**Supplementary Figure S1. CD55 is highly expressed on CSCs. (A)** CSC and non-CSC histograms for additional membrane-bound complement inhibitory proteins, CD59 and CD46. **(B)** mRNA expression was determined by qPCR and compared between GFP+ (CSCs) and GFP- (non-CSCs) enriched from A2780 cells using the NANOG-GFP reporter system. Actin was used as a control. **(C)** CSCs were also enriched by surface CD49f expression in A2780, which demonstrated higher CD55 levels at protein and mRNA levels. **(D, E, F)** Cisplatin-naïve and –resistant CSCs vs non-CSCs histogram plots for CD55 expression. **(G)** Cell lysates from cisplatin-resistant CSCs and non-CSCs were immunoblotted and probed for CD55. Actin was used as a loading control. **(H)** Limiting dilution analysis plots of CD55+ and CD55- cisplatin resistant cells. The graph compares the estimates of the percentage of self-renewal frequency in these sorted populations with the corresponding p-values. *p< 0.5, **p< 0.01, ***p< 0.001

**Supplementary Figure S2. CD55 maintains self-renewal in cisplatin-resistant endometrioid tumors. (A)** Immunoblots of cisplatin-naïve A2780 CSCs with CD55 silencing and non-targeted control were probed with CD55, CD59, and CD46. Actin was used as loading control. **(B)** mRNA expression was determined by qPCR and compared between CD55-silenced CSCs and non-targeted control CSCs. Actin was used as a control. **(C)** Immunoblots of cisplatin-resistant parental cells silenced for CD55 using two shRNA constructs and a non-targeting control were probed with CD55, NANOG, SOX2, and OCT4. Actin was used as a loading control. **(D)** Limiting dilution analysis plots of CD55 NT control compared with CD55 sh1 and sh2 silencing constructs in cisplatin-resistant parental cells. **(E)** Cell lysates from CD59 silenced A2780 CSCs and their non-targeted (NT) controls were immunoblotted and probed with CD59, NANOG, SOX2, and OCT4. Actin was used as loading control. **(F)** Limiting dilution analysis plots of CD59 NT control compared with CD59 KD1 and KD2 silencing constructs in A2780 CSCs. *p< 0.5, **p< 0.01, ***p< 0.001

**Supplementary Figure S3. CD55 maintains platinum resistance in patient-derived xenograft and cisplatin-resistant endometrioid tumors. (A)** CD55 silenced cisplatin-naïve uterine PDX CSCs and their NT controls were treated with cisplatin, percentage of surviving cells and relative caspase 3/7 activity were graphed. (B) Relative caspase 3/7 activity for CD55 silenced cisplatin-naïve cells and their NT controls after treatment with cisplatin. <u>Data are representative of three independent experiments.</u> **(C, D)** CD55 silenced cisplatin-resistant parental cells and their NT controls were treated with cisplatin, percentage of surviving cells and relative caspase 3/7 activity were graphed. **(E)** In vivo cisplatin sensitivity studies were performed comparing the NT control and CD55-silenced group. Graph shows the growth rate of tumors compared to the first day of cisplatin treatment. **(F)** CD59 silenced A2780 CSCs and their NT controls were treated with 0-50 μM cisplatin and percent surviving cells are graphed. *p< 0.5, **p< 0.01, ***p< 0.001

**Supplementary Figure S4. CD55 regulates self-renewal and cisplatin resistance in a complement independent manner. (A)** Complement-mediated cytotoxicity was assessed by the percentage of BCECF dye release in CSCs vs non-CSCs, and cisplatin resistant vs naïve cells treated with 10, 20, and 30% normal human serum (NHS). **(B)** A2780 cells sorted based on their surface CD55 expression were treated with 10 and 20% NHS, and growth relative to untreated controls was graphed. **(C)** Limiting dilution analysis plots of CD55+ and CD55- A2780 cells cultured with or without 10% NHS. **(D)** CD55+ and CD55- A2780 cells cultured with or without NHS were treated with 0-50 μM cisplatin and percent surviving cells were graphed. **(E)** Immunofluorescent staining of cisplatin-naïve CSCs was performed for CD55 and cholera toxin B. **(F)** PIPLCtreated CSCs and their vehicle-treated controls were treated with 0-50 μM cisplatin and percent surviving cells are graphed. **(G)** Receptor tyrosine kinase array was performed against 71 unique tyrosine kinases to identify the pathways altered by CD55 silencing in CSCs. **(H)** CSCs of cisplatin-naïve cells were sorted based on their surface CD55 expression and immunoblotted for CD55, ROR2, pLCK (Y394), and LCK. Actin was used as loading control. *p< 0.5, **p< 0.01, ***p< 0.001. Scale bar: 4 μm.

**Supplementary Figure S5. CD55 signals via LCK and induces DNA repair genes. (A)** Cell lysates from TOV112D CSCs and non-CSCs were immunoblotted and probed with pLCK (Y394) and LCK. Actin was used as loading control. **(B)** Pull-down experiments with CD55 antibody were performed in CP70 parental cells and elutes were probed for pLCK (Y394), LCK, and CD55. **(C)** Limiting dilution analysis plots of saracatinib and DMSO-treated cisplatin-naïve CSCs. **(D)** Cell lysates from LCK silenced A2780 CSCs and their non-targeted (NT) controls were immunoblotted and probed with LCK. Actin was used as loading control. **(E)** LCK silenced CSCs and their NT controls were treated with 0-50 μM cisplatin and percent surviving cells are graphed. **(F)** Limiting dilution analysis plots of cisplatin-naïve non-CSCs transduced with LCK overexpression and empty vector constructs. **(G)** mRNA expression was determined by qPCR and compared between LCKoverexpressing non-CSCs and empty vector control non-CSCs. Actin was used as a control. **(H)** Growth curves for CD55-overexpressing non-CSCs and their empty vector controls treated with or without saracatinib. The graph shows growth relative to day 0. **(I)** mRNA fold changes of the four most significantly modulated genes from gene expression profiling, comparing non-CSCs transduced with empty vector control vs CD55 or LCK overexpression, and CSCs with non-targeted control vs CD55-silencing, and CSCs with DMSO vs saracatinib treatment. **(J, K)** MLH1 and BRCA1 silenced CSCs and their non-targeted controls were treated with 0-50 μM cisplatin and percent surviving cells are graphed. ^‡^Emtpy vector control for non-CSCs, and non-targeted control for CSCs. *p< 0.5, **p< 0.01, ***p< 0.001

## REFERENCES

Armstrong DK, Bundy B, Wenzel L, Huang HQ, Baergen R, Lele S, Copeland LJ, Walker JL, Burger RA, Gynecologic Oncology G. 2006. Intraperitoneal cisplatin and paclitaxel in ovarian cancer. N Eng J Med 354:34–43.

Baldwin LA, Huang B, Miller RW, Tucker T, Goodrich ST, Podzielinski I, DeSimone CP, Ueland FR, van Nagell JR, Seamon LG. 2012. Ten-year relative survival for epithelial ovarian cancer. Obstet Gynecol 120:612–618.

Cancer Genome Atlas Research Network., K.C., Schultz N, Cherniack AD, Akbani R, Liu Y, Shen H, Robertson AG, Pashtan I, Shen R, Benz CC, Yau C, Laird PW, Ding L, Zhang W, Mills GB, Kucherlapati R, Mardis ER, Levine DA. 2013. Integrated genomic characterization of endometrial carcinoma. Nature 497:67–73.

Catasus L, Gallardo A, Cuatrecasas M, Prat J. 2009. Concomitant PI3K-AKT and p53 alterations in endometrial carcinomas are associated with poor prognosis. Mod Pathol 22:522–529.

CGAR, N. 2011. Integrated genomic analyses of ovarian carcinoma. Nature 474:609–615.

Cuellar-Partida G, Lu Y, Dixon SC; Australian Ovarian Cancer Study., Fasching PA, Hein A, Burghaus S, Beckmann MW, Lambrechts D, Van Nieuwenhuysen E, Vergote I, Vanderstichele A, Doherty JA, Rossing MA, Chang-Claude J, Rudolph A, Wang-Gohrke S, Goodman MT, Bogdanova N, Dörk T, Dürst M, Hillemanns P, Runnebaum IB, Antonenkova N, Butzow R, Leminen A, Nevanlinna H, Pelttari LM, Edwards RP, Kelley JL, Modugno F, Moysich KB, Ness RB, Cannioto R, Høgdall E, Høgdall C, Jensen A, Giles GG, Bruinsma F, Kjaer SK, Hildebrandt MA, Liang D, Lu KH, Wu X, Bisogna M, Dao F, Levine DA, Cramer DW, Terry KL, Tworoger SS, Stampfer M, Missmer S, Bjorge L, Salvesen HB, Kopperud RK, Bischof K, Aben KK, Kiemeney LA, Massuger LF, Brooks-Wilson A, Olson SH, McGuire V, Rothstein JH, Sieh W, Whittemore AS, Cook LS, Le ND, Blake Gilks C, Gronwald J, Jakubowska A, Lubiński J, Kluz T, Song H, Tyrer JP, Wentzensen N, Brinton L, Trabert B, Lissowska J, McLaughlin JR, Narod SA, Phelan C, Anton-Culver H, Ziogas A, Eccles D, Campbell I, Gayther SA, Gentry-Maharaj A, Menon U, Ramus SJ, Wu AH, Dansonka-Mieszkowska A, Kupryjanczyk J, Timorek A, Szafron L, Cunningham JM, Fridley BL, Winham SJ, Bandera EV, Poole EM, Morgan TK, Goode EL, Schildkraut JM, Pearce CL, Berchuck A, Pharoah PD, Webb PM, Chenevix-Trench G, Risch HA, MacGregor S. 2016. Assessing the genetic architecture of epithelial ovarian cancer histological subtypes. Hum Genet 135:741–756.

DiSaia PJ, Creasman W. 2012. Clinical Gynecologic Oncology. Elsevier, Philadelphia.

Galluzzi L, Vitale I., Michels J, Brenner C, Szabadkai G5 Harel-Bellan A, Castedo M, Kroemer G. 2014. Systems biology of cisplatin resistance: past, present and future. Cell Death Dis 5:e1257.

Gyorffy B, Lanczky A, Szallasi Z. 2012. Implementing an online tool for genome-wide validation of survival-associated biomarkers in ovarian-cancer using microarray data of 1287 patients. Endocr Relat Cancer 19:197–208.

Hanker LC, Loibl S, Buchrdi N, Pfisterer J, Meier W, Pujade-Lauraine E, Ray-Coquard I, Sehouli J, Harter P, du Bois A, Ago, group Gs. 2012. The impact of second to sixth line therapy on survival of relapsed ovarian cancer after primary taxane/platinum-based therapy. Ann Oncol 23:2605–2612.

Horejsí V, Zhang W, Schraven B. 2004. Transmembrane adaptor proteins: organizers of immunoreceptor signalling. Nat Rev Immunol 4:603–616.

Hu Y, Smyth GK. 2009. ELDA: extreme limiting dilution analysis for comparing depleted and enriched populations in stem cell and other assays. J Immunol Methods 347:70–78.

Ikeda J, Morii E, Liu Y, Qiu Y, Nakamichi N, Jokoji R, Miyoshi Y, Noguchi S, Aozasa K. 2008. Prognostic significance of CD55 expression in breast cancer. Clin Cancer Res 14:4780–4786.

Kapka-Skrzypczak L, Wollinska E, Szparecki G, Czajka M, Skrzypczak M. 2015. The immunohistochemical analysis of membrane-bound CD55, CD59 and fluid-phase FH and FH-like complement inhibitors in cancers of ovary and corpus uteri origin. Centr Eur J Immunol 40:349–353.

Karst AM, Drapkin R. 2010. Ovarian cancer pathogenesis: a model in evolution. J Oncol 2010:932371.

Kurman RJ, Shih I. 2016. The Dualistic Model of Ovarian Carcinogenesis: Revisited, Revised, and Expanded. Am J Pathol 186:733–747.

Kyo S, Maida Y, Inoue M. 2011. Stem cells in endometrium and endometrial cancer: accumulating evidence and unresolved questions. Cancer Lett 308:123–133.

Lathia JD, Gallagher J, Heddleston JM, Wang J, Eyler CE, Macswords J, Wu Q, Vasanji A, McLendon RE, Hjelmeland AB, Rich JN. 2010. Integrin alpha 6 regulates glioblastoma stem cells. Cell Stem Cell 6:421–432.

Lathia JD, Li M., Sinyuk M, Alvarado AG, Flavahan WA, Stoltz K, Rosager AM, Hale J, Hitomi M, Gallagher J, Wu Q, Martin J, Vidal JG, Nakano I, Dahlrot RH, Hansen S, McLendon RE, Sloan AE, Bao S, Hjelmeland AB, Carson CT, Naik UP, Kristensen B, Rich JN. 2014. High-throughput flow cytometry screening reveals a role for junctional adhesion molecule a as a cancer stem cell maintenance factor. Cell Rep 6:117–129.

Li Y, Lin F. 2012. Mesenchymal stem cells are injured by complement after their contact with serum. Blood 120:3436–3443.

Lukacik P, Roversi P., White J, Esser D, Smith GP, Billington J, Williams PA, Rudd PM, Wormald MR, Harvey DJ, Crispin MD, Radcliffe CM, Dwek RA, Evans DJ, Morgan BP, Smith RA, Lea SM. 2004. Complement regulation at the molecular level: the structure of decay-accelerating factor. Proc Natl Acad Sci U S A. 101:1279–1284.

Murray KP, Mathure S, Kaul R, Khan S, Carson LF, Twiggs LB, Martens MG, Kaul A. 2000. Expression of complement regulatory proteins-CD 35, CD 46, CD 55, and CD 59-in benign and malignant endometrial tissue. Gynecol Oncol 76:176–182.

Nagaraj AB, Joseph P, Kovalenko O, Singh S, Armstrong A, Redline R, Resnick K, Zanotti K, Waggoner S, DiFeo A. 2015. Critical role of Wnt/β-catenin signaling in driving epithelial ovarian cancer platinum resistance. Oncotarget 6:23720–23734.

Oishi, I., H. Suzuki, N. Onishi, R. Takada, S. Kani, B. Ohkawara, I. Koshida, K. Suzuki, G. Yamada, G.C. Schwabe, S. Mundlos, H. Shibuya, S. Takada, and Y. Minami. 2003. The receptor tyrosine kinase Ror2 is involved in non-canonical Wnt5a/JNK signalling pathway. Genes Cells 8:645–654.

Qiu, C., Y. Ma, J. Wang, S. Peng, and Y. Huang. 2010. Lin28-mediated post-transcriptional regulation of Oct4 expression in human embryonic stem cells. Nucleic Acids Res 38:1240–1248.

Saxe, J.P., Tomilin, A., H.R. Scholer, K. Plath, and J. Huang. 2009. Post-translational regulation of Oct4 transcriptional activity. PLoS One 4:e4467.

Shenoy-Scaria AM, Kwong J, Fujita T, Olszowy MW, Shaw AS, Lublin DM. 1992. Signal transduction through decay-accelerating factor. Interaction of glycosyl-phosphatidylinositol anchor and protein tyrosine kinases p56lck and p59fyn 1. J Immunol 149:3535–3541.

Siegel RL, Miller KD., Jemal A. 2016. Cancer statistics, 2016. CA Cancer J Clin 66:7–30.

Tan TZ, M.Q., Huang RY, Wong MK, Ye J, Lau JA, Wu MC, Bin Abdul Hadi LH, Soong R, Choolani M, Davidson B, Nesland JM, Wang LZ, Matsumura N, Mandai M, Konishi I, Goh BC, Chang JT, Thiery JP, Mori S. 2013. Functional genomics identifies five distinct molecular subtypes with clinical relevance and pathways for growth control in epithelial ovarian cancer. EMBO Mol Med 5:1051–1066.

Thiagarajan PS, Hitomi M, Hale JS, Alvarado AG, Otvos B, Sinyuk M, Stoltz K, Wiechert A, Mulkearns-Hubert E, Jarrar AM, Zheng Q, Thomas D, Egelhoff TT, Rich JN, Liu H, Lathia JD, Reizes O. 2015. Development of a Fluorescent Reporter System to Delineate Cancer Stem Cells in Triple-Negative Breast Cancer. Stem Cells 33:2114–2125.

Unno K, Ono M, Winder AD, Maniar KP, Paintal AS, Yu Y, Wei JJ, Lurain JR, Kim JJ. 2014. Establishment of human patient-derived endometrial cancer xenografts in NOD scid gamma mice for the study of invasion and metastasis. PLos One 9:e116064.

Ventimiglia LN, Alonso MA. 2013. The role of membrane rafts in Lck transport, regulation, and signaling in T-cells. Biochem J 454:169–179.

Wiechert A, Saygin C, Thiagarajan PS, Rao VS, Hale JS, Gupta N, Hitomi M, Nagaraj AB, DiFeo A, Lathia JD, Reizes O. 2016. Cisplatin induces stemness in ovarian cancer. Oncotarget 7:30511–30522.

Yao, Y., Lu, Y., W.C. Chen, Y. Jiang, T. Cheng, Y. Ma, L. Lu, and W. Dai. 2014. Cobalt and nickel stabilize stem cell transcription factor OCT4 through modulating its sumoylation and ubiquitination. PLoS One 9:e86620.

